# H2AK119ub Safeguards Against Ectopic Transcription Factor Mediated Gene Activation in the Developing Forebrain

**DOI:** 10.1101/2025.07.07.663591

**Authors:** Lucy A. Doyle, Alexandra Derrien, Dipta Sengupta, Firuze Unlu Bektas, Jennifer Lawson, Chinmayi Pednekar, Alexander Von Kriegsheim, Ian R. Adams, Robert S. Illingworth

## Abstract

The PRC1 complex regulates developmental gene expression in mammals by ubiquitinating histones (H2AK119ub) and nucleating repressive chromatin interactions. Human genetic data and functional experimentation have demonstrated that H2AK119ub is required for cortical development, however the molecular mechanism for this remains unknown. Here, we show that mouse embryos expressing catalytically deficient RING1B exhibit intact early neurogenesis but display impaired neuroectodermal fate restriction. Using in vivo, primary, and in vitro models, we demonstrate that reduced H2AK119ub leads to ectopic activation of lineage-inappropriate transcriptional programmes, including mesodermal and endodermal gene expression during neural differentiation. This transcriptional deregulation is not solely attributable to H2AK119ub or H3K27me3 loss but instead reflects sensitisation of PRC1 target genes to inappropriate transcription factor (TF)-mediated activation. Synthetic induction of candidate TFs, including GATA6, SOX7, and SNAI1, phenocopies the fate-skewing effects of PRC1 catalytic dysfunction, confirming their causal role. Our results uncover a buffering role for PRC1-catalysed H2AK119ub in safeguarding neural progenitor identity by preventing inappropriate TF-driven transcription and provides a mechanistic framework for understanding the cellular heterogeneity and phenotypic variability observed in Polycomb-associated neurodevelopmental disorders.

## Introduction

Polycomb group (PcG) proteins are essential regulators of mammalian development. Integrated into multi-subunit complexes, PcGs control gene expression by remodelling the chemical and physical properties of chromatin^1-3^. Polycomb repressive complex 1 (PRC1) and polycomb repressive deubiquitinase (PR-DUB), act antagonistically to control the levels of the repressive histone mark - H2A lysine 119 monoubiquitination (H2AK119ub). PRC1 deposits ubiquitin through the action of a core catalytic heterodimer comprising the E3 ubiquitin ligase RING1A or B and one of six paralogous PCGF subunits^4-6^. In contrast, PR-DUB removes H2AK119ub from chromatin by cleaving ubiquitin from the H2A tail, via the action of the C-terminal hydrolase BAP1^7-9^. PRC1 and PR-DUB therefore act in combination to control the level and spatial distribution of H2AK119ub within the genome^10-13^. Non-canonical (nc)PRC1 complexes containing either RYBP or YAF2 are responsible for the majority of H2AK119ub deposition, whereas canonical (c)PRC1 acts primarily to nucleate chromatin interactions^6,14-16^.

The molecular mechanism by which H2AK119ub represses gene expression remains unclear, however recent evidence suggests that it operates by reducing the frequency of transcriptional bursting^17,18^. To explore the extent to which H2AK119ub shapes global transcription levels, we and others engineered mouse ESCs that encode catalytically deficient forms of RING1B (*Ring1b^I53A^*, *Ring1b^I53S^* and *Ring1b^I53A/D56K^*). Each of these mutations disrupts the interaction between RING1B and the E2 ubiquitin-conjugating enzyme, whilst preserving its incorporation into otherwise functional PRC1 complexes^19-24^. These mutations, either alone or in combination with deletion of *Ring1a*, cause varying degrees of H2AK119ub loss (80-100% global and 50-100% focal reduction at PRC1-target sites). An 80% global (50-60% focal) loss of H2AK119ub causes only modest gene mis-regulation, however a more substantial reduction leads to failure of PRC1-mediated repression^20,22,24^. This suggests that there is a minimum level of H2AK119ub required to support PRC1-mediated repression in mouse ESCs.

Mammalian embryogenesis produces specialized tissues though progressive rounds of cellular proliferation, differentiation, migration, and spatial patterning. Development of the neocortex (corticogenesis), the brain region responsible for higher cognitive functions, follows these same basic principles^25, 26, 27^. Multipotent neural stem cells (NSCs) undergo sequential and temporally constrained rounds of expansion, neurogenesis and gliogenesis^25,28^. During neurogenesis, new-born neurons populate deep ventricle-proximal regions, with subsequent-rounds migrating to increasingly superficial positions culminating in the laminar arrangement that is characteristic of the cortex^25-27,29^. The timing of these transitions, the regulation of cell fate and the spatial organization of migrating cells are all essential to ensure the correct size and structure of the cortex^30-34^. Corticogenesis is regulated by interconnected cell-intrinsic and extrinsic mechanisms that include extracellular signalling, gene regulatory networks and hierarchical cascades of transcription factors (TFs) ^29,33-37^. These mechanisms converge on chromatin to establish the appropriate cell-type specific gene expression patterns. However, corticogenesis is a highly dynamic process and relies upon the PcG system to facilitate flexible repression of master TFs^28,38-44^.

The importance of H2AK119ub during corticogenesis is underscored by the existence of monogenic neurodevelopmental disorders (NDDs) that are caused by disruption of the H2AK119ub regulatory axis (reviewed in ^45-47^). Mutations in both PRC1 and PR-DUB give rise to phenotypically related NDDs, suggesting that an ‘optimal’ level of H2AK119ub1 is required for appropriate gene regulation and normal brain development^48-55^. In experimental models, loss of RING1B in primary NSCs extends the neurogenic window and delays the onset of gliogenesis due to an inability to silence pro-neural gene such as *Neurog1*^41^. Separation of function analysis shows that this repression is dependent upon H2AK119ub in early neurogenic NSCs but that it is dispensable for repression of the same genes in gliogenic NSCs. This suggests that H2AK119ub supports a ‘primed’ repressive state that can then be removed or replaced by more stable repressive mechanisms in a context specific manner^44^. However, it is not clear mechanistically how primed H2AK119Ub-containing chromatin impacts on gene regulation to promote neuronal development. Here we investigate the molecular and developmental impact of reduced H2AK119ub deposition in the embryonic mouse cortex, primary NSCs and in vitro derived neural precursor cells (NPCs). Using these systems, we investigate the key molecular and cellular sensitivities of impaired H2AK119ub-deposition. Our findings show that PRC1-mediated H2AK119ub operates at the functional interface between chromatin and TFs to safeguard neuroectodermal development.

## RESULTS

### NSCs derived from *Ring1b^I53A/I53A^* mice exhibit reduced neuroectodermal fate restriction

The catalytic activity of RING1B is required for the repression of pro-differentiation genes in neural progenitor cells and therefore to control the timing of brain development^41,44,56^. To determine the impact of reduced H2AK119ub deposition on neurodevelopment in vivo, we employed a mouse model that constitutively expresses a catalytically deficient form of RING1B (*Ring1b^I53A/I53A^*) ^20^. Unlike RING1B knockouts, homozygous embryos survive to mid-gestation at approximately Mendelian frequency, with loss of embryos occurring only at later developmental stages (≥ E15.5) ^20^. Importantly, this allowed us to investigate the impact of reduced H2AK119ub deposition on neurogenesis (E12.5), a critical window during which NSCs expand and differentiate into neuronal fated progenitors and neurons.

At a gross scale, *Ring1b^I53A/I53A^* embryos were smaller (∼10% reduction; crown-to-rump length and head width) with a disproportionately greater reduction in the size of the forebrain (∼20% reduction; **Supplementary** Fig. 1a, b). Visual inspection of coronal sections revealed a reduction in the thickness of both dorsal (cerebral cortex) and ventral structures (lateral and medial ganglionic eminences - LGE and MGE; **Fig. 1a**) in *Ring1b^I53A/I53A^* embryos. Despite this, immunofluorescent detection of SOX2 (stem cell marker) and TUJ1 (neuron-specific β-Tubulin) showed stage-appropriate distribution of sub/ventricular zone (SVZ/VZ) NSCs and cortical plate (CP) neurons. This indicated that the reduced size of the forebrain did not reflect a substantial developmental delay or an inability to produce cortical neurons in *Ring1b^I53A/I53A^* embryos (**Fig. 1a, b** and **Supplementary** Fig. 1a, b).

**Fig. 1:**
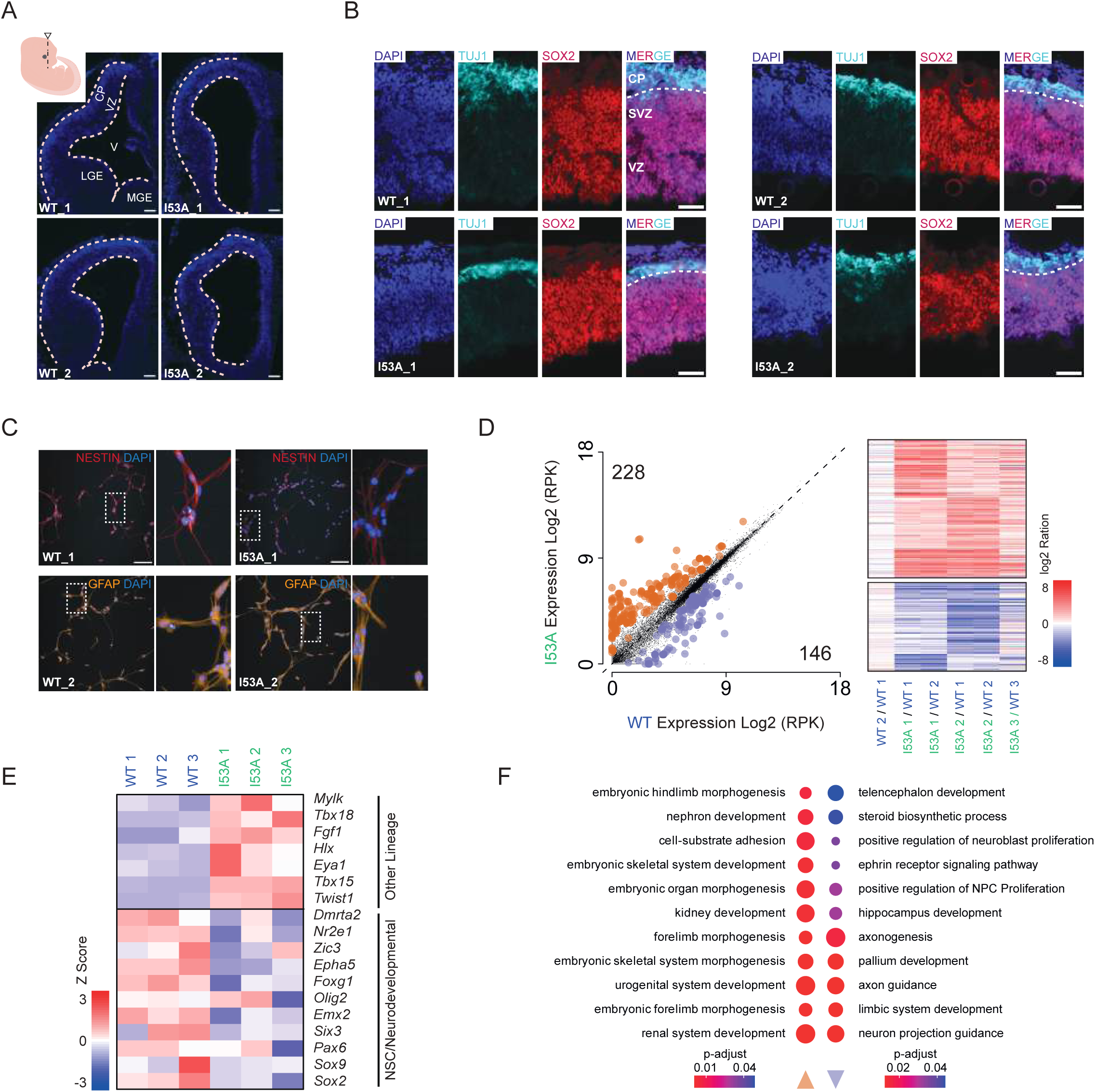
Reduced neuroectodermal fate restriction in primary NSCs derived from *Ring1b^I53A/I53A^* mice. **a,** Representative DAPI stained coronal forebrain sections of *Ring1b^+/+^* (WT) and *Ring1b^I53A/I53^*(I53A) E12.5 mouse embryos (section plane indicated in inset cartoon schematic). Cortical and ventricular surfaces are outlined and scale bars = 100 μm. **b,** Representative fluorescent images of DAPI staining and TUJ1 (β-tubulin) and SOX2 immunofluorescence of *Ring1b^+/+^* (WT) and *Ring1b^I53A/I53A^* (I53A) E12.5 coronal cortical sections (scale bar = 50 μm). **a** and **b**, LGE = lateral ganglionic eminence, MGE = medial ganglionic eminence, V= ventricle, VZ = ventricular zone, SVZ = sub-ventricular zone and CP = cortical plate. **c,** Representative fluorescent images of DAPI staining in combination with either NESTIN or GFAP immunofluorescence in *ex vivo* cultured primary telencephalic NSCs derived from *Ring1b^+/+^* (WT) and *Ring1b^I53A/I53A^*(I53A) E10.5 mouse embryos (scale bar = 100 μm; right hand panel shows a zoomed view as indicated). **d,** (Left panel) Scatter plots comparing mean normalised gene expression (Log_2_ normalised read per kilobase - RPK) between *Ring1b^+/+^*(WT) and *Ring1b^I53A/I53A^* (I53A) NSCs. Differentially expressed genes (DEGs) which are significantly up- or downregulated are indicated in red and blue respectively (log_2_ FC ≥ 1 / ≤ -1 and p ≤ 0.01) and the numbers of DEGs are indicated. (Right panel) Heatmaps depicting the log_2_ FC (log2 I53A/WT) for upregulated and downregulated genes for independent NSC cultures derived from two litters of embryos (litter 1 - WT 1, WT 2, I53A 1 and I53A 2; litter 2 - WT 3 and I53A 3). The heatmap data range is scaled as per the colour bar. **e,** Heatmap comparing the gene expression profiles (z-score) of candidate developmental genes in NSCs cultured from *Ring1b^+/+^* (WT) and *Ring1b^I53A/I53A^* (I53A) embryos. The heatmap is scaled as per the colour bar. **f,** The top eleven enriched functional gene ontology terms for upregulated and downregulated DEGs shown in **e**. Colour indicates the adjusted p value as per the colour bar and the circle size indicates the gene count per GO term.

We reasoned that whilst new-born neurons at the cortical plate were being produced in *Ring1b^I53A/I53A^*embryos (**Fig. 1b**), appropriate levels of H2AK119ub may be required to safeguard the transcriptional and developmental fidelity of the progenitor cells that give rise to them. To test this, we isolated primary radial progenitor (RP) NSCs directly from the telencephalons of *Ring1B^I53A/I53A^* mutants^57^. Cultured NSCs from wild-type and *Ring1B^I53A/I53A^* E10.5 embryos presented with the characteristic bi-polar morphology of RPs and stained uniformly positive for both NESTIN and GFAP by immunofluorescence (**Fig. 1c** and **Supplementary** Fig. 1c, d). RNA-seq performed on independent litter matched *Ring1B^I53A/I53A^* and *wildtype* NSCs, identified a total of 374 differentially expressed genes (DEGs), the majority of which (228/374 ∼61%) were upregulated in *Ring1B^I53A/I53A^* NSCs (**Fig. 1d; Supplementary Table 1**). These expression changes were consistent across NSCs cultured from independent mouse litters (**Fig. 1d**; Litter 1 – WT 1, 2 and I53A 1, 2 and Litter 2 – WT 3 and I53A 3). Inspection of individual DEGs revealed downregulation of core forebrain NSC/neurodevelopmental marker genes (e.g. *Sox2*, *Pax6*, *Emx2* and *Foxg1*) and upregulation of marker genes more commonly associated with other cell lineages (e.g. *Twist1 and Tbx15;* **Fig. 1e** and **Supplementary** Fig. 1e). Consistent with this, functional gene ontology (GO) analysis showed ectopic upregulation of markers of the neural crest, mesoderm and endoderm and a reciprocal reduction in the expression of genes encoding pro-neural regulators (**Fig. 1f; Supplementary Table 2**). These findings demonstrate that RING1B catalytic activity is required to safeguard neuroectodermal fate restriction in NSCs that were established within their normal *in vivo* developmental niche.

### Transcription factor motif enrichment and not H2AK11ub loss defines genes that are upregulated in *Ring1b^I53A/I53A^* NSCs

Upregulation of a subset of non-neuroectodermal genes in NSCs, highlighted a gene-selective sensitivity to impaired RING1B-function (**Fig. 1d; Supplementary Table 1**). We reasoned therefore, that although the NSC genome showed a global reduction in H2AK119ub levels of approximately 35%, variable losses across TSSs could sensitise specific genes to ectopic activation (**Fig. 2a-c** and **Supplementary** Fig. 2a). Consistent with this notion, calibrated chromatin immunoprecipitation (cChIP-seq) showed that upregulated genes had on average, higher levels of TSS proximal H2AK119ub in WT NSCs, and a more pronounced reduction in the levels of the modification in *Ring1b^I53A/I53A^* mutants (**Fig. 2d, e, Supplementary** Fig. 2b and **Supplementary Table 3**). However, whilst this appeared to be necessary for activation, it was not sufficient, as many genes with equivalent H2AK119ub dynamics were not upregulated (**Fig. 2f, Supplementary** Fig. 2c and **Supplementary Table 3)**.

**Fig. 2:**
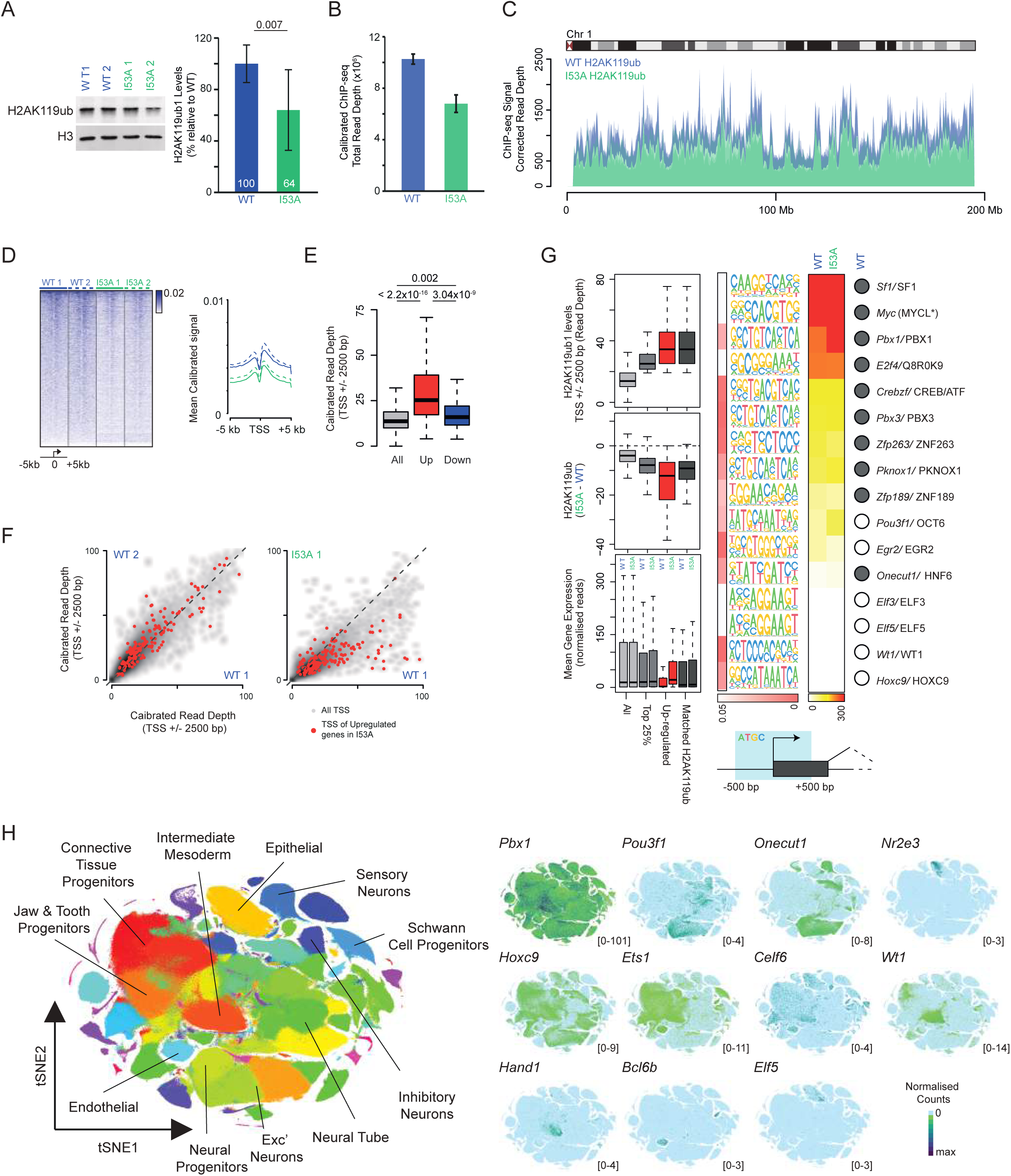
TF motif enrichment and not the extent of H2AK11ub loss defines genes that are upregulated in *Ring1b^I53A/I53A^* NSCs. **a,** (left) Immunoblot of histone H2AK119ub and histone H3 and (right) a barplot showing the mean level of H2AK119ub from two independent *Ring1b^+/+^*and *Ring1b^I53A/I53A^* primary NSC cultures. The relative levels (percentage of WT) and the significance (determined using a Welch Two Sample t-test) are indicated. Error bars represent the standard deviation of eight replicates (2 biological and 4 technical per genotype). **b,** Barplot showing the mean normalised uniquely mapping read depth following calibrated ChIP-seq for H2AK119ub1 in two independent *Ring1b^+/+^* and *Ring1b^I53A/I53A^* NSC cultures. **c,** Calibrated ChIP-seq profiles for H2AK119ub1 in *Ring1b^+/+^* and *Ring1b^I53A/I53A^* (WT and I53A; blue and green profiles respectively) for mouse chromosome 1 (mm39 genome build). Tracks for 2 independent biological and 2 technical replicates are shown for each genotype. **d,** Heatmaps (left panel) and summary profiles (right panel) of calibrated H2AK119ub ChIP-seq signal prepared from the indicated NSC extracts. Heatmaps are ranked from high to low signal based on the WT 1 NSC sample (mean of two technical replicate experiments). Profiles display the ChIP-seq signal surrounding the TSSs (±5 kb). **e,** Boxplots of wildtype H2AK119Ub levels at the TSS (±2.5 kb) of all (grey), upregulated in *Ring1b^I53A/I53A^* (red) and downregulated in *Ring1b^I53A/I53A^* (blue) NSCs. Significant changes between datasets (determined using a Wilcoxon rank sum test) are indicated above their respective plot. **f,** Scatter plots showing the H2AK119ub1 levels at TSSs (±2.5 kb) in the indicated NSC cultures. H2AK119ub values for TSSs of genes that are upregulated in *Ring1b^I53A/I53A^* NSCs are highlighted in red. **g,** (Left panel) Boxplots showing absolute H2AK119ub1 levels (TSS ±2.5 kb) in *Ring1b^+/+^* NSCs, differential H2AK119ub1 levels (TSS ±2.5 kb) in *Ring1b^I53A/I53A^* - *Ring1b^+/+^* NSCs and gene expression in *Ring1b^+/+^* and *Ring1b^I53A/I53A^* NSCs (top, middle and bottom respectively). Normalised ChIP-seq and RNA-seq values are shown for the indicated gene sets. (Central panel) Significantly enriched binding motifs found in the TSSs (±500 bp – see cartoon schematic) of genes that are upregulated in *Ring1b^I53A/I53A^* vs. *Ring1b^+/+^* NSCs. Adjusted p-value are indicated in the red gradient heatmap. (Right panel) Heatmap showing RNA levels for the genes encoding the TFs (normalised RPK values scaled as per the colour bar). Filled circles indicate detectable protein expression in both *Ring1b^+/+^* NSCs determined using Mass Spectrometry. **h,** tSNE plots depicting normalised scRNA-seq data from the ‘Mouse Organogenesis Cell Atlas’ generated using the UCSC single cell browser ^59,103^. (Left panel) tSNE plot depicting dimensionally reduced expression data coloured based the major cell type as defined by selected maker gene expression. Selected relevant cell types are indicated for reference. (Right panel) tSNE plots showing the cell-type-specific expression patterns of a selection of TFs whose motifs are enriched in the H2AK119ub positive *Ring1b ^I53A/I53A^*NSC upregulated genes characterised in **g**. Normalised expression range is indicated for each gene in parenthesis and coloured according to the colour bar.

To determine if alterations in PcG function more generally underpin gene selectivity we performed calibrated ChIP-seq for the PRC2 mark H3K27me3 in *Ring1b^+/+^* and *Ring1B^I53A/I53A^* NSCs (**Supplementary** Fig. 2a). TSSs with high WT levels of H3K27me3 did show a reduction in *Ring1B^I53A/I53A^*NSCs and, as for H2AK119ub, this effect was more pronounced at upregulated genes (**Supplementary** Fig. 2d, e and **Supplementary Table 3**). Indeed, both H3K27me3 and H2AK119ub profiles revealed a high level of correlation when comparing the relative loss of the two modifications at individual gene loci demonstrating that, as for ESCs, H3K27me3 deposition in NSCs is partially dependent on PRC1-mediated H2AK119ub^20,58^. However, pairwise comparison of differential H2AK119ub and H3K27me3 levels at gene TSS, showed that whilst upregulated genes had a more substantial reduction in both modifications, many genes whose expression was unchanged showed an equivalent loss (**Supplementary** Fig. 2f and **Supplementary Table 3**). This demonstrated that a reduction in PRC-associated modifications, either alone or in combination, could not account for the observed gene-specific activation.

We speculated then, that H2AK119ub could be acting to restrict the binding or action of context-specific sequence dependent transcription factors (TFs), and that reduced PRC1 activity could therefore make certain genes more susceptible to ectopic TF-mediated activation. To test this, we identified genes with high levels of H2AK119ub proximal to their TSS (top 25%) and that were either upregulated in *Ring1B^I53A/I53A^*NSCs, or a randomly selected matched set whose expression did not change (**Fig. 2g -** left panel). Transcription factor (TF) motif enrichment analysis comparing these two sets identified 30 significantly enriched motifs in the TSSs of the upregulated, H2AK119ub positive genes (**Fig. 2g** – middle panel; **Supplementary** Fig. 2g – middle panel and **Supplementary Table 4** and **5**). We found that whilst many of these TFs were expressed broadly across embryonic cell types (e.g. *Pbx1* and *Hoxc9*), the subset of TFs with a more restricted developmental profile were generally enriched in non-neuroectodermal lineages (e.g. *Wt1*, *Ets1* and *Hand1;* **Fig. 2h**) ^59^. These findings suggest that loss of H2AK119ub in NSCs sensitise PRC1-target genes to inappropriate activation by non-neuroectodermal driver TFs.

### Reduced H2AK11ub levels causes ectopic activation of endodermal and mesodermal genes during neuroectodermal specification

The transition from stem cell to neuron is molecularly highly challenging and our analysis captured a ‘snapshot’ only, and did not allow the impact of reduced H2AK119ub to be determined as neuroectodermal specification occurs. To address this, we investigated the capacity of *Ring1B^I53A/I53A^* mouse ESC to differentiate into neural progenitors and neurons^20^. Immunoblotting of histone preparations from *Ring1B^I53A/I53A^ mESCs* showed a greater than five-fold reduction in global H2AK119ub levels relative to *Ring1B^+/+^* controls (**Fig. 3a**) ^20,22^. This loss was more substantial than that observed in primary NSCs (**Fig 2a-c**), likely due to a higher rate of cell division and reduced compensation by RING1A. We induced differentiation of *Ring1B^I53A/I53A^* mESCs toward a neural fate by withdrawing leukaemia inhibitory factor (LIF) and culturing them in neural specification conditions (**Supplementary** Fig. 3a) ^20,60,61^. Following 14-days of differentiation, both *Ring1B^+/+^* and *Ring1B^I53A/I53A^*cells presented with characteristic TUJ1 positive neurite outgrowths (**Fig. 3b, c** and **Supplementary** Fig. 3b, c). Notably however, *Ring1B^I53A/I53A^* cultures contained “cobble-stone” shaped cells and large clusters of cells with spontaneous contractile properties that resembled cardiomyocytes, a phenomenon that was consistent across replicate experiments, but not observed in *Ring1B^+/+^* controls (**Supplementary Movie 1-3**).

**Fig. 3:**
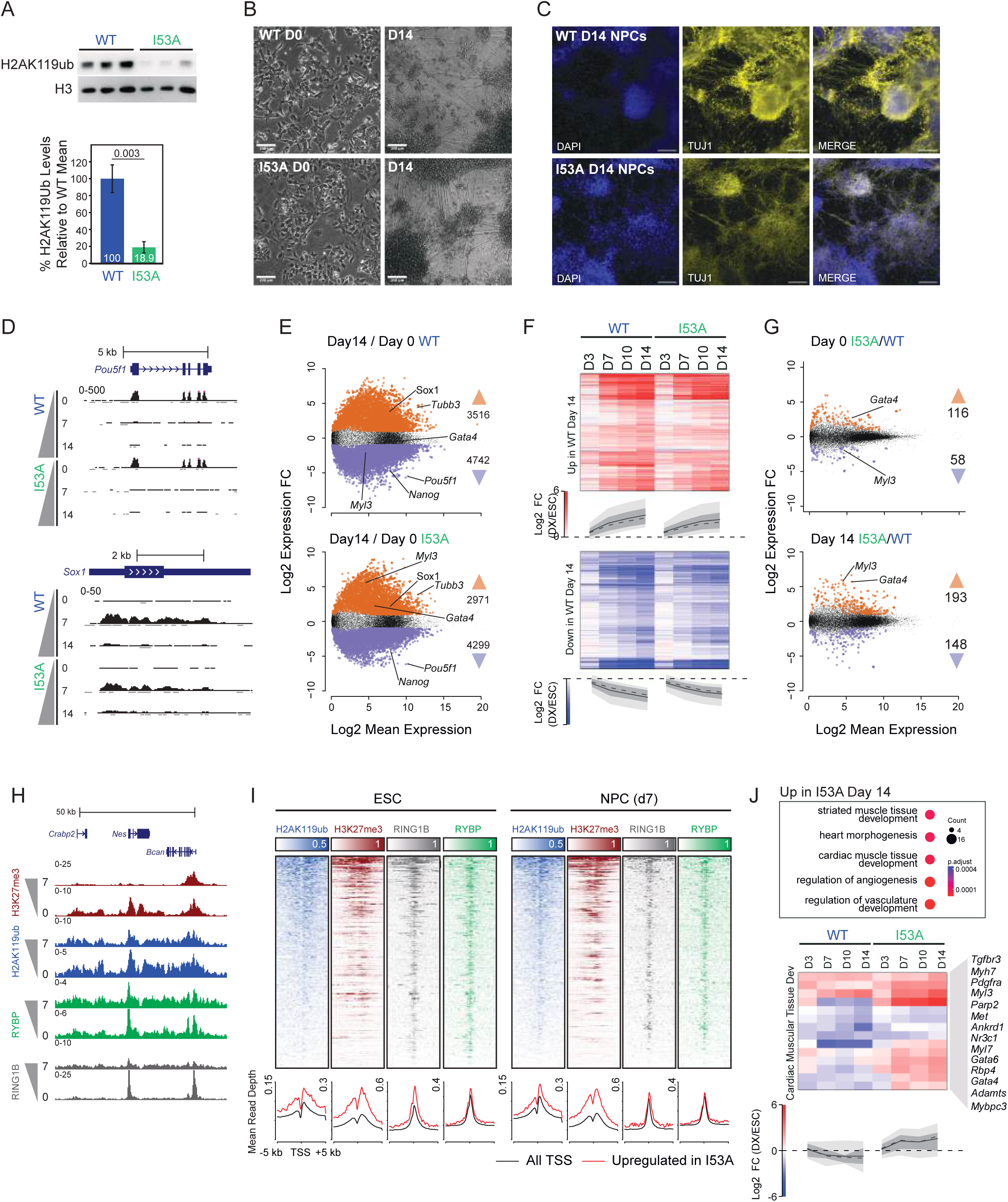
*Ring1b^I53A/I53A^*ESCs display non-lineage appropriate cell specification during directed NPC differentiation. **a,** Immunoblot of histone H2AK119ub and histone H3 (upper panel) and barplot showing the relative levels of H2AK119ub (lower panel) in *Ring1b^+/+^* and *Ring1b^I53A/I53A^* mESCs. The relative levels (percentage of WT) and the p value (determined using a Welch Two Sample t-test) are indicated. Error bars show the standard deviation of three independent replicates. **b,** Representative bright-field images of *Ring1b^+/+^* and *Ring1b^I53A/I53A^* cells at day 0 and 14 of NPC differentiation (scale bar = 100 μm). **c,** Representative fluorescent images of DAPI staining and TUJ1 (β-tubulin) immunofluorescence in *Ring1b^+/+^* and *Ring1b^I53A/I53A^*cells at day 14 of NPC differentiation (Scale bar = 100 μm). **d,** Genome browser tracks of normalised RNA-seq signal at candidate gene loci (*Pou5f1 and Sox1*) at day 0, 7 and 14 of NPC differentiation for *Ring1b^+/+^* (upper) and *Ring1b^I53A/I53A^* (lower) cells. The signal is coloured according to the transcribed strand (positive - black and negative - grey) and annotated as per the mm39 genome assembly. The scale of normalised read-depth is indicated. **e,** Scatter plots comparing log_2_ mean expression vs. log_2_ fold changes (FC; log2 Day 14/Day 0) for *Ring1b^+/+^*and *Ring1b^I53A/I53A^* cells at day 14 of NPC differentiation. Differentially expressed genes (DEGs) which are significantly up- or downregulated are indicated in red and blue respectively (log_2_ FC ≥ 1 / ≤ -1 and p ≤ 0.01). Candidate marker genes and the total number of DEGs are indicated for each comparison. **f,** Heatmaps depicting the log_2_ fold changes (FC; log2 Day X/Day 0) for upregulated and downregulated genes (upper and lower panel respectively) are shown with their combined meta-profiles (dashed line = median, solid line = mean, dark grey = 25^th^ -75^th^ percentile and light grey 10^th^ – 90^th^ percentile). The heatmap is scaled as per the axis colour bar. **g,** Scatter plots comparing log_2_ mean expression vs. log_2_ FC (I53A/WT) at day 0 and 14 of NPC differentiation (displayed as for **e**). **h,** Genome browser tracks of ChIP-seq signal for H3K27me3, H2AK119ub, RYBP and RING1B at candidate genomic loci (*Nes*) in WT ESCs (‘0’) and their derivative NPCs (‘7’). The regions are annotated as per the mm39 genome assembly and normalised read-depth range is indicated. **i,** Heatmaps of H3K27me3, H2AK119ub, RYBP and RING1B ChIP-seq signal spanning ±5 kb of the TSSs of upregulated genes in ESCs and NPCs (left and right panels). Heatmaps are ranked from high to low signal based on the respective H2AK119ub datasets. Profiles plots are shown below each heatmap respective heatmap depicting the modification profile at upregulated genes (red) relative to all genes (black). **j,** The top five enriched functional gene ontology terms for upregulated DEGs and heatmap of log_2_ FC (I53A/WT) for the indicated functional gene list. The heatmaps and profiles are shown as for **f**.

To investigate the molecular basis of this phenotype, we performed RNA-seq across the differentiation, sampling at days 0 (ESCs), 3, 7, 10 and 14 (**Supplementary** Fig. 3a). Inspection of normalised reads at candidate loci showed a reduction in expression of the pluripotency factor *Pou5f1* (the gene encoding OCT4) and a reciprocal upregulation of the early neural marker *Sox1* as differentiation progressed in both *Ring1B^+/+^* and *Ring1B^I53A/I53A^*cells (**Fig. 3d**). Consistent with this observation, and the presence of TUJ1+ cells in later stage cultures, global analysis showed a broadly similar gene expression profile between the two genotypes (**Fig. 3e, f, Supplementary** Fig. 3d and **Supplementary Table 6**). However, we also noted upregulation of non-lineage appropriate genes such as the master endodermal regulator *Gata4* and the cardiac myosin *Myl3* in *Ring1B^I53A/I53A^*cells (**Supplementary** Fig. 3d, e and **Supplementary Table 6**). Expression of genes that are not usually active in ectodermal cells prompted us to directly compare the two genotypes throughout the differentiation. By day 14, 341 genes were differentially expressed, the majority of which (193/341) were upregulated in *Ring1B^I53A/I53A^*cells, consistent with our findings in primary NSCs (**Fig. 1d, Fig. 3g, Supplementary** Fig. 3f-h and **Supplementary Table 6**). To determine if this was due to direct up-regulation of PcG targets, we performed ChIP-seq for RING1B and RYBP (PRC1 subunits), H2AK119ub (PRC1 modification) and H3K27me3 (PRC2 modification) in *Ring1b^+/+^*mESCs and d7 NPCs (**Fig. 3h** and **Supplementary** Fig. 3i). Quantification of normalised ChIP-seq signal showed that upregulated gene sets had on average, higher levels of both marks and their mediators in ESCs and NPCs, confirming that they were enriched for direct targets of PcG-mediated repression (**Fig. 3i** and **Supplementary Table 7**).

We next wanted to determine if the observed gene mis-regulation was indicative of skewed fate determination as indicated by ectopic *Gata4* expression. GO analysis identified a significant enrichment for functions involved in mesodermal and endodermal development, including ‘striatal and cardiac muscle development’, ‘heart morphogenesis’ and ‘regulation of vasculature development’, in genes upregulated in *Ring1B^I53A/I53A^*differentiation (**Fig. 3j** and **Supplementary Table 8**). This came at a cost to neural lineage specification with downregulated genes being enriched for functions involved in ‘forebrain neuron differentiation’ and ‘CNS neuronal differentiation’ (**Supplementary** Fig. 3j and **Supplementary Table 8**). These trends are illustrated in the quantitative reduction of key neural markers and the reciprocal increased expression of endodermal and mesodermal markers in differentiated *Ring1B^I53A/I53A^* cells (**Fig. 3j, Supplementary** Fig. 3j and **Supplementary Table 8**).

This skew in developmental potential was similar to that observed in primary NSCs (**Fig. 1**). To compare the two models without the confounder of cellular heterogeneity, we cultured *Ring1B^+/+^* and *Ring1B^I53A/I53A^* ESCs, first in NPC differentiation media, and then in NSC media containing EGF and FGF^57^. The resulting homogeneous population morphologically resembled primary NSCs and clustered with them based on their global gene expression profile determined by RNA-seq (**Supplementary** Fig. 1c, **Supplementary** Fig. 4a-d and **Supplementary Table 9**). Differential gene expression analysis between *Ring1B^I53A/I53A^*and *Ring1B^+/+^* in vitro derived NSCs identified 1,048 DEGs (**Supplementary** Fig. 4e and **Supplementary Table 9**). The differentially expressed genes overlapped significantly with those identified in primary NSCs, with the most substantial overlap being between downregulated gene sets (**Supplementary** Fig. 4f, g). Functional GO analysis showed that this overlap reflected a clear and consistent loss of neuroectodermal fate restriction in both primary and in vitro derived *Ring1B^I53A/I53A^* NSCs (**Supplementary** Fig. 4h and **Supplementary Table 10**). Our data suggests that PRC1 catalytic activity is required to suppress mesoderm and endodermal gene expression programmes and restrict specification to neural lineages during early development.

### Activation of key transcription factors phenocopies the impact of H2AK119ub loss in NPCs

Our results suggest that reduced H2AK119ub levels cause the erosion of the transcriptional identity of neural progenitor cells by increasing their sensitivity to low levels of lineage inappropriate TFs. Analysis of the TF encoding genes which are differentially expressed between WT and *Ring1B^I53A/I53A^* identified two distinct clusters; those which are upregulated early only and those which progressively increase during the course of NPC differentiation (category 1 and 2 respectively; **Fig. 4a** and **Supplementary Table 6**). As the emergence of beating heart cells occurred late in NPC differentiation, we primarily focussed on the TFs encoded by category 2 genes (**Supplementary Movies 1-3**). Of these, we identified SOX7, GATA6 and SNAI1 as putative drivers of the observed phenotype, as they collectively contribute to mesodermal, endodermal and cardiac development, as well as inhibiting ectodermal gene expression in embryonic mesoderm^62-67^.

**Fig. 4:**
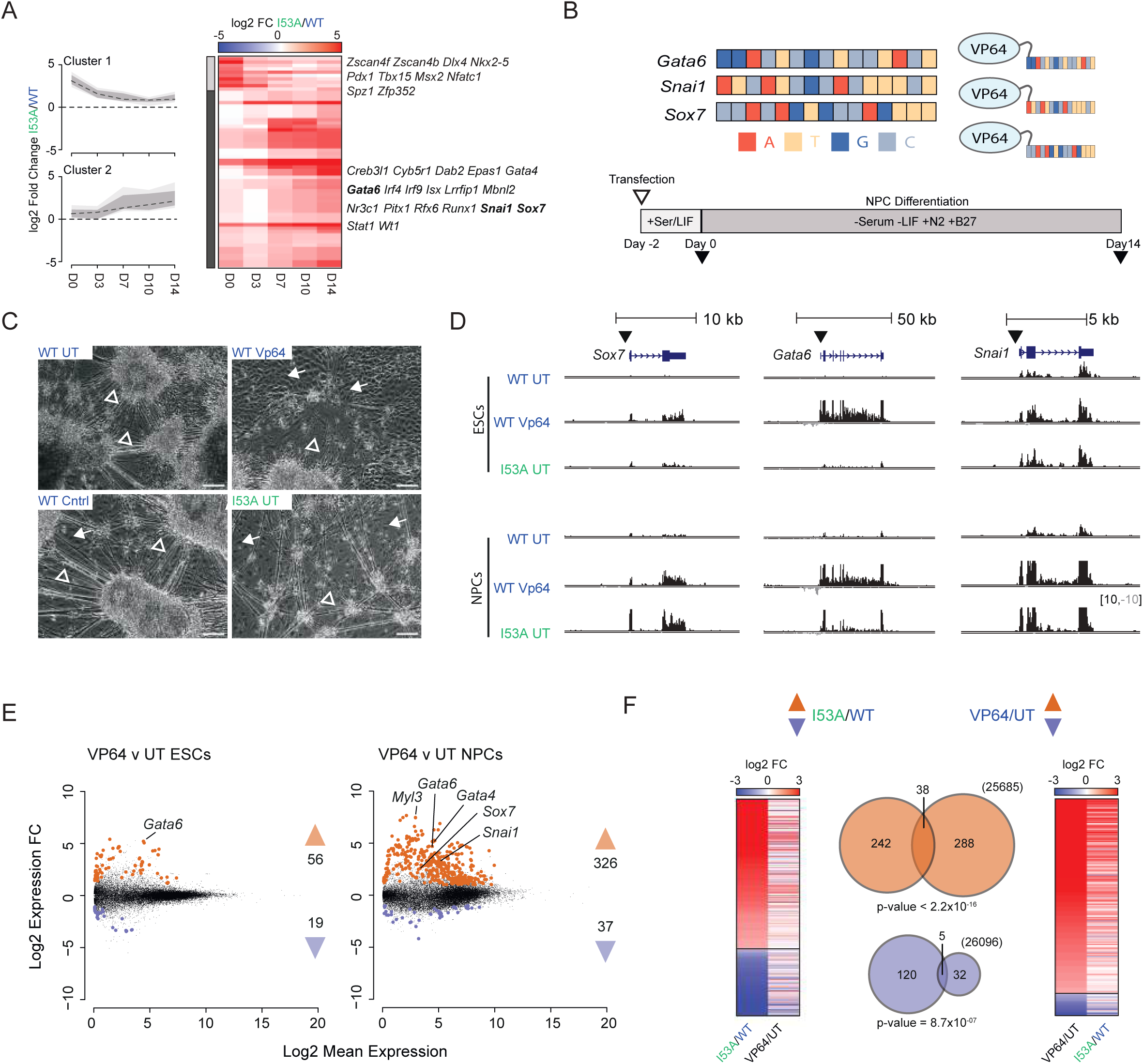
Activation of master developmental TFs phenocopies the transcriptional impact of H2AK119ub loss during NPC differentiation. **a,** Summary profiles and heatmap depicting the log_2_ fold changes (I53A/WT) for TF encoding genes that are upregulated in I53A relative to WT cells during NPC differentiation. Genes were separated into two groups by hierarchical clustering, and the summary profiles show the expression dynamics of each cluster (dashed line = median, dark grey = 25^th^ -75^th^ percentile and light grey 10^th^ – 90^th^ percentile). The heatmap is scaled as per the axis colour bar and the TF genes for each cluster are shown on the right of the heatmap. **b,** Cartoon schematic of the TALE-VP64 constructs and the differentiation strategy. Each TALE repeat variable domain (RVD) monomer is coloured according to its target DNA base. **c,** Representative bright-field images of un-transfected WT cells (WT UT), WT transfected with TALE-delta control (WT Cntrl), WT transfected with TALE-VP64 (WT VP64) and I53A un-transfected control (I53A UT) following 14 days under NPC differentiation conditions. Open and filled arrows indicate neurite projections and cobble-stone-like cells respectively (scale bar = 100 μm). **d,** Genome browser tracks of mean normalised RNA-seq signal at target gene loci (*Sox7, Gata6* and *Snai1*) in ESCs and NPCs (day 14) for the indicated conditions. The signal represents the sum of three independent experimental replicates and is coloured according to the transcribed strand (positive - black and negative - grey). Regions are annotated as per the mm39 genome assembly. Filled arrows indicate the target site for TALE constructs in relevant samples. **e,** Scatter plots of log_2_ mean gene expression vs. log_2_ fold change (FC) for WT cells for the indicated comparisons at day 0 (ESCs; left) and 14 (NPCs; right) of differentiation. Differentially expressed genes (DEGs) that are significantly up- or downregulated are indicated in red and blue respectively (log_2_ FC ≥ 1 / ≤ -1 and p ≤ 0.01). Candidate marker genes and the total number of DEGs are indicated for each comparison. **f,** Heatmaps and matched Venn Diagrams comparing DEGs identified in I53A vs WT NPCs (day 14; left panel) and VP64 vs. UT Control cells (right panel). The significance of the overlap for both sets of genes was determined using a Fisher’s Test, the result of which is shown below each respective Venn.

If *Ring1B^I53A/I53A^* cells respond inappropriately to low levels of these TFs, we reasoned that the developmental constraint imposed by high levels of H2AK119ub in WT cells could be overcome by elevating the expression of these same TFs. To test this, we employed a synthetic approach to transiently activate the three TFs in ESCs and monitor the impact of this on the outcome of NPC differentiation. We cloned transcription activator-like effector (TALE) constructs encoding a targeting module (recognising 16 base pairs proximal to the TSS of each TF gene) fused to the potent synthetic activator VP64 (**Fig. 4b**). We introduced these constructs as a pool into WT ESCs and cultured them in NPC differentiation media for 14 days. Whilst ESCs showed no overt phenotype, 14-day cultures transfected with the VP64-fusion pool displayed a reduction in the numbers of cells with neurite projections and the emergence of a population of “cobble-stone” shaped cells; morphology that resembled that of un-transfected *Ring1B^I53A/I53A^* cells cultured under matched conditions (**Fig. 4c** and **Supplementary** Fig. 5a, b). In contrast, WT cells transfected with targeting constructs lacking a VP64 cargo (‘Cntrl’) were indistinguishable from UT ESCs, indicating that activation of the TFs directly underpinned the apparent changes in cell composition (**Fig. 4c** and **Supplementary** Fig. 5a, b).

As enforced expression of GATA6, SOX7 and SNAI1 in WT ESCs altered the outcome of targeted NPC differentiation in a manner that resembled *Ring1B^I53A/I53A^* cells, we sought to determine if this was underpinned by changes in gene expression. Visual inspection of RNA-seq signal from ESCs and NPCs from each condition confirmed the activation of the target genes, and that this persisted in NPCs at a level equivalent to that in *Ring1B^I53A/I53A^* cultures (**Fig. 4d** and **Supplementary Table 11**). Differential expression analysis comparing activated cells (‘VP64’) with either un-transfected cells (‘UT’) or cells transfected with the control (‘Cntrl’) identified a substantial shift in gene expression in Day 14 NPCs (363 and 311 DEGs in D14 NPCs respectively; **Fig. 4e, Supplementary** Fig. 5c and **Supplementary Table 11**). Differentially expressed genes between *Ring1B^I53A/I53A^* and WT NPCs overlapped significantly with those identified between VP64 and UT NPCs, particularly for upregulated genes (**Fig. 4f** and **Supplementary Table 11**). This is consistent with the observation that both *Gata4* and *Myl3*, genes that were found to be differentially expressed between *Ring1B^I53A/I53A^* and WT NPCs and known targets of the selected TFs, are upregulated upon TF induction (**Fig. 4e, Supplementary** Fig. 4c, d and **Supplementary Table 11**). The phenocopying of the transcriptional response shows that GATA6, SOX7 and SNAI1 directly contribute to the *Ring1B^I53A/I53A^*differentiation phenotype. This demonstrates that RING1B-mediated H2AK119ub safeguards neurodevelopmental fidelity by buffering against inappropriate TF-mediated activation.

## DISCUSSION

PRC1 is an essential regulator of mammalian brain development that contributes both catalytic and structural chromatin modifying functions to control NSC expansion and differentiation ^40,41,44,56,68,69^. In this study we demonstrate that mouse embryos that express a catalytically deficient form of RING1B (*Ring1B^I53A/I53A^*) produce NSCs and early born neurons. However, in primary NSCs, the reduction in H2AK119ub leads to impaired neuroectodermal fate restriction and ectopic activation of genes typically associated with alternative cell lineages. In vitro, *Ring1B^I53A/I53A^*ESCs produced beating cardiomyocyte-like cells under NPC specifying conditions, providing strong physiological evidence of an aberrant cell fate switch (**Fig. 5a**). This transition was driven by activation of a subset of PRC1 target genes by sequence specific TFs including SOX7, SNAI1 and GATA6. Therefore, a combination of global de-repression and target-site selective activation defines the transcriptional and developmental outcomes of impaired RING1B-catalysis.

**Fig. 5:**
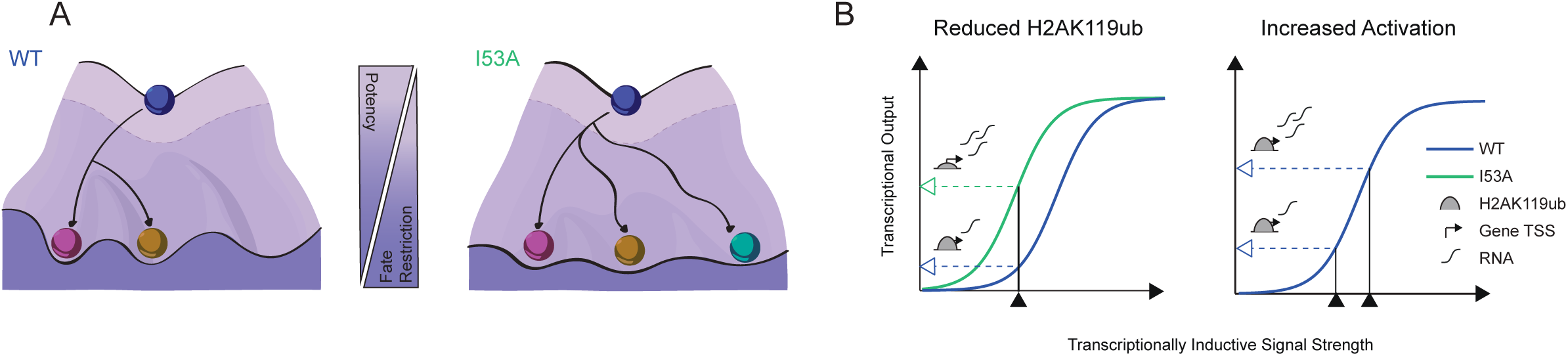
Altering the balance between H2AK119ub levels and inductive developmental TFs de-rails neuroectodermal specification. **a,** Cartoon depiction styled on Conrad Waddington’s valley model indicating the developmental consequence of impaired RING1B function (*Ring1b^I53A/I53A^*; I53A) on neuroectodermal specification. **b,** Model schematic depicting how both reduced levels of TSS-associated H2AK119ub (left) or increasing the level of an activating TF (right) leads to increased expression of target genes.

### The Impact of PRC1 Catalytic Dysfunction is Dictated by Cellular Context

To determine the influence of developmental constraint on PRC1-mediated gene regulation, we investigated the consequence of the *Ring1b^I53A/I53A^* mutation in mouse embryos, primary cultured NSCs and ESC-derived NPCs. The erroneous emergence of mesodermal cells in vitro, is in stark contrast to the seemingly subtle structural and cytological perturbations observed in embryos bearing the same mutation. Furthermore, both primary cultured and in vitro derived *Ring1b^I53A/I53A^*NSCs displayed reduced neuroectodermal fate restriction but presented with more distinct patterns of gene upregulation (**Supplementary** Fig. 4). Our model that reduced levels of global H2AK119ub lowers the threshold for ectopic TF-mediated gene activation (**Fig. 5b**) provides an explanation for this context specificity, as TFs driving this response will be cell-type-specific. Impaired PRC1 function will therefore allow a given cell to deviate from its developmental trajectory, but the manner and extent to which it does so will be dictated and constrained by its developmental context. Indeed, progenitor cells established in vivo will be more developmentally constrained by the rich signalling and mechanical cues of their niche environment. This is reflected in the frequent occurrence of developmental growth abnormalities observed in PcG-associated chromatinopathies, where altered cell abundance indicates skewed developmental timing, rather than wholesale developmental failure^45-47^. This mechanistic insight is important for understanding how altered levels of H2AK119ub could drive disease in different developmental and cellular contexts.

### Imbalanced H2AK119ub in NDD Chromatinopathies

Despite their opposing activities, mutations in both PRC1 and PR-DUB give rise to phenotypically overlapping NDDs suggesting that an ‘optimal’ level of H2AK119ub1 is required for typical brain development^48-55^. While PR-DUB removes H2AK119ub to prevent inappropriate repression, its loss paradoxically results in upregulation of a subset of PRC1-target genes^7,10-13^. This reflects a dual role for the H2AK119ub writer-eraser axis; both facilitating landmark deposition at target genes and preventing inappropriate accumulation elsewhere. Disruption of this balance leads to misallocation of limited PRC resources, promoting both ectopic gene repression and target gene de-repression^11,12^. The erosion of neurodevelopmental fidelity observed in this study resembles that caused by loss of PCGF6 (PRC1.6) in ESC derived NPCs, supporting the idea that impaired H2AK119ub deposition disrupts neuroectodermal specification^6,70^. It is unclear whether individual mutations in PRC1 and PR-DUB converge on a unified transcriptional signature, however disruption of the balance achieved by the ‘monoubiquitin axis’ could represent a common aetiology.

### H2AK119ub and Functional Crosstalk Between PRCs

The contribution of H2AK119ub to gene regulation is increasingly understood as part of a dynamic interaction with other PcG components^22,24,71^. In ESCs, canonical PRC1 recruitment is facilitated by PRC2-mediated H3K27me3 via CBX7, while PRC2 is stabilized on chromatin by H2AK119ub through AEBP2 and JARID2^58,72-78^. We show that reduced H2AK119ub in mutant NSCs results in decreased H3K27me3 at PRC1 target loci, most prominently at upregulated genes (**Supplementary** Fig. 2c-f). Whilst this effect may be amplified by displacement of PRC2 from chromatin due to association with nascent RNA ^79,80^, reduced H3K27me3 levels were also observed at the TSSs of genes with no detectible change in expression (**Supplementary** Fig. 2c-f). Combined with the fact that AEBP2, JARID2 and CBX7 are expressed at appreciable levels in NSCs, this suggests that the reciprocal mechanisms for PRC recruitment also exists in NSCs (**Supplementary Table 6**). What contribution this reduction in H3K27me3 (PRC2) levels has on the observed transcriptional upregulation is unknown, but this finding highlights the importance of H2AK119ub in preserving overall PRC occupancy in diverse cell-types.

### Concluding remarks

Our findings provide strong evidence that RING1B-catalysis plays a central role in controlling cell-fate in neural progenitors; a finding that is consistent with the phenotypes caused by disruption of H2AK119ub regulation. In cases such as the recently described mis-sense mutations in RING1B^81^, loss-of-function is predicted to have genome-wide consequences. Yet brain development in affected individuals is grossly typical, suggesting that the developmental gene expression programme is largely intact. That many genes are not-misregulated even upon loss or absence of H2AK119ub suggests that targeted gene activation is also required in addition to global de-repression. A better understanding of how specific gene-expression signatures result from such derepression is critical to understanding the molecular basis of polycomb-associated NDDs. Furthermore, dysfunction of other PRC components and indeed other chromatin modifiers such as MBD3/NuRD also loosen the reins on lineage choice^44,70,82-85^. Understanding the interplay between chromatin modifiers will guide development of therapeutic strategies aimed at correcting or mitigating chromatinopathy phenotypes^86^.

## METHODS

### Ring1B^I53A/I53A^ mice

Generation of *Ring1B^I53A/I53A^*mice was described previously^20^. Homozygotes were produced through heterozygote matings (*Ring1B^+/I53A^* x *Ring1B^+/I53A^*cross). For embryonic staging, the detection of a vaginal plugs in mated Dams was taken as embryonic day E0.5. Pregnant females were culled by cervical dislocation and E10.5 and E12.5 embryos were removed and transferred to ice cold 1x PBS and stored on wet ice for 0.5 – 3 h. A small portion of the tail tip was surgically removed at this point and genotyped as previously described^20^. Analysis of *Ring1B^I53A/I53A^* mice was performed under a UK Home Office project license (PPL 60/4424) with approval from an institutional ethics committee.

### Cell Culture and Differentiation

Feeder-free E14TG2a and the derivative line *Ring1B^I53A/I53A^* ^20^ were cultured as previously described ^23^.

NPC differentiation was performed essentially as described in ^60^. Briefly, ESCs were seeded at high density (6 × 10^4^ cells/cm^2^) on 0.1% gelatin-coated flasks (Corning) and cultured in ESC media for 24 h. ESCs were harvested and washed first with room temperature (RT) 1x PBS and then RT 1x NPC base media (1:1 mixture of DMEM/F-12 (Gibco – 31330-038) and Neurobasal media (Gibco - 21103049)). Washed ESCs were seeded on 0.1% gelatin-coated 6-well culture plates (Corning) at a density of approximately 1×10^4^ cells/cm^2^ in NPC differentiation media (NPC base media supplemented with 1x B27 (Gibco - 17504044), 1x N-2 (Gibco - 17502048), 0.2 mM L-glutamine (Gibco - 25030123) and 0.1 mM 2-Mercaptoethanol (Gibco - 31350010)) and cultured for 7 – 14 days without passaging. Cells were harvested either by trypsinisation or by mechanical scraping.

Primary NSCs from E10.5 embryos were cultured essentially as previously described ^57^. Briefly, the telencephalons of E10.5 embryos were surgically removed and mechanically disaggregated in RT 1x PBS (Gibco -10010023) by pipetting with a wide bore pipette tip. Single cells were retrieved by passing the cells through a 40 µm cell strainer (Greiner - 542040) and collected by centrifugation. NSCs were re-suspended in complete media (DMEM/F-12 (Gibco – 31330-038), 1:100 Penicillin-Streptomycin (Pen/Strep; Gibco 15140-122), 7.7 mM glucose (Sigma G8644), 1X non-essential amino acids (Gibco - 11140035), 0.01% BSA (Gibco; 15260-037), 0.2 mM L-glutamine (Gibco - 25030123), 0.1 mM 2-Mercaptoethanol (Gibco - 31350010), 1x B27 (Gibco - 17504044), 1x N-2 (Gibco - 17502048), 10 ng/ml mouse EGF (Peprotech - 315-09), 10 ng/ml human FGF (Peprotech - 100-18b), 2 µg/ml Cultrex laminin (Trevigen; 3446-005-01)) and were seeded in a single well of a 6-well plate (Corning). For passaging, 60-90% confluent wells were washed with RT 1x PBS, incubated for 2–3 min at 37 °C in 1x Accutase (133 µl/cm^2^ surface area; Thermo-Fisher - 00-4555-56), and tapped to release. The cells were diluted in 5x volumes of wash media (DMEM/F-12, 1:100 Penicillin-Streptomycin, 7.7 mM glucose, 1X non-essential amino acids, 0.01% BSA and 0.2 mM L-glutamine). NSCs were centrifuged, resuspended in complete media and replated onto laminin coated wells at a ratio of between 1:4 and 1:8. Where necessary, improved cellular adherence was achieved by pre-treating culture plates with 5 µg/ml Laminin in 1x PBS for 20 - 30 min at RT or overnight at 4 °C.

Unless otherwise stated, all centrifugation steps involving live cells were performed at 330 x g for 3 min at RT. Cells were cultured in a humidified incubator at 37 °C with 5% CO_2_.

### Histone Extraction and Immunoblotting

Histones were isolated as previously described^87^ and then quantified by BCA assay (according to manufacturer’s instructions; Pierce - 23225). All samples were equalised to 1 µg in 1x Sample Loading Buffer (31 mM Tris-HCl pH 6.8, 12.5 % Glycerol, 0.005 % (w/v) Bromophenol Blue, 1 % w/v SDS and 100 mM DTT) and denatured at 90 °C for 3 min. Samples were vortexed and centrifuged for 5 min at 14,000 x g to clear the lysates and then stored at −20°C until required. Samples were loaded on to a hand-cast 15% SDS-PAGE gel and resolved by electrophoresis in 1x Tris Glycine Running Buffer (25 mM Tris base, 192 mM glycine and 0.1% w/v SDS) at 100 V for 5 min and then 200 V for 70 min. Resolved protein was transferred to nitrocellulose membrane (0.2 µm; Biorad - 1620112) by wet transfer (Mini-PROTEAN Tetra Blotting Module; Biorad - 1660827EDU) at 100 V for 45 min in chilled Transfer Buffer (25 mM Tris base, 190 mM Glycine and 20% w/v Methanol). After two washes in 1x PBS, membranes were blocked in PBST (1x PBS and 0.1 % v/v Tween 20) supplemented with 10% w/v skimmed milk powder and incubated ON at 4 °C on a shaker. Following two washes in 1x PBS, membranes were incubated with primary antibody in Antibody Buffer (1x PBST supplemented with 50mg/ml BSA) for 3h at 4°C with agitation. Membranes were washed 5 x 5 min in RT PBST and then incubated with secondary antibody diluted in PBST for 1h at RT with agitation. H2AK119ub and pan-H3 were probed using Rabbit anti-H2AK119ub D27C4 (1:2000; Cell Signalling - 8240S) and Mouse anti-Histone H3 96C10 (1:1000; Cell Signalling - 3638S) and then labelled using Goat Anti-Rabbit IgG Alexa Fluor 647 (1:2000; Thermo Fisher - A-21244) and Goat Anti-Mouse IgG Alexa Fluor 488 (1:2000; Thermo Fisher - A-11001) respectively. All antibody solutions were centrifuged at >14,000 x g for 15 min prior to use to remove antibody aggregates. Membranes were washed 4 x 5 min in RT PBST and 1 x 5 min in 1 x PBS. Blots were imaged on a ChemiDoc MP Imaging System (Biorad) and quantified using the Fiji software package^88^. To ensure comparable densitometry results, gel electrophoresis, protein transfer and antibody staining were performed in parallel for all samples.

### Immunofluorescent Microscopy

#### Sample Preparation

E12.5 Embryos were fixed overnight in 4% paraformaldehyde (PFA) at 4 °C, rinsed in RT 1x PBS, then passed through increasing concentrations of sucrose in 1x PBS (10%, 20% and 30% w/v), each for >4 h at 4 °C. Once equilibrated, the tissue was embedded in OCT (CellPath - KMA-0100-00A), orientated using forceps and frozen by placing the casting mould on a bed of dry ice, and then stored at -70 °C. Tissue was cryo-sectioned into 10 µm sections using a cryostat (Leica), mounted on Superfrost plus slides (VWR – 631-0108) and then stored at - 70°C. Cells were cultured on glass coverslips pre-treated with 15% v/v Poly-L-Ornithine (Sigma - P4957) in 1x PBS overnight at 4 °C and then with either 10 µg/ml Laminin in 1x PBS for 3-6 h at 4 °C for NSCs or 0.1% gelatin in 1x PBS for 3-6 h at 4 °C for ESC to NPC differentiations. Cells were fixed for 10 min at RT in 4% PFA and stored at -20 °C.

#### Immunofluorescent detection

Slides/coverslips were washed 3 x 5 min in RT 1 x PBS. Antigen retrieval was performed on sections by microwaving slides for 10 min in sodium citrate buffer (10 mM Sodium citrate pH 6.0 and 0.05% Tween20,). A barrier perimeter was drawn around each sample using a hydrophobic barrier pen (Vector Laboratories – H4000) to retain the block and hybridisation mixtures used in subsequent steps. Samples were permeabilised in Blocking Buffer (10% sheep serum and 0.1% v/v Triton X100 (Sigma-Aldrich - 102152669)) for 1 h at RT. Primary antibodies (see below) were diluted in Blocking Buffer and incubated overnight at 4°C (a negative control omitting the primary antibody was also included). Slides/coverslips were washed 3 x 5 min with RT 1 x PBS, and then probed with fluorescent secondary antibodies diluted to 1:250 in 1 x PBS for 1 h at RT. Slides/coverslips were washed 3 x 5 min with RT 1 x PBS, and air dried at RT protected from light. Samples were mounted with ProLong Gold Antifade fluorescent mounting media (Thermo Fisher - P36930), sealed with nail varnish and stored at 4 °C. Immunofluorescent imaging was carried out using a Zeiss Observer microscope and analysed using the Fiji software package^88^. β-tubulin (TUBB3), NESTIN, GFAP and SOX2 were probed using Mouse anti-TUJ1 (1:400; BioLegend - 801201), Mouse anti-NESTIN (1:20; DSHB - rat-401), Chicken anti-GFAP (1:500; BioLegend - 829401) and Rabbit anti-SOX2 (1:200; Abcam - ab92494) and then labelled using Donkey anti-Mouse IgG Alexa Fluor 647 (1:250; Thermo Fisher - A-31571), Goat anti-Chicken IgY Alexa Fluor Plus 488 (1:250; Thermo Fisher – A32931) and Donkey anti-Rabbit IgG Alexa Fluor 568 (1:250; Thermo Fisher – A10042) respectively.

### Protein Expression Analysis by Mass Spectrometry

Harvested cells were washed with 1 x PBS and pelleted by centrifugation at 300 x g for 5 min. Proteins were prepared by incubation in Lysis Buffer (6 M Gu-HCl, 1 mg/ml Chloroacetic acid (CAA; alkylates free cysteines) and 1.5 mg/ml tris(2-carboxyethyl)phosphine (TCEP; reduces disulphide bonds)) at 95 °C for 10 min. Proteins were then double digested using Lys-C and Trypsin (Promega). Once digested, the peptides were desalted by loading on to C18 hydrophobic membrane stage tips and then quantified using a NanoDrop spectrophotometer (Thermo Fisher).

Desalted peptides were loaded onto 25 cm Aurora Columns (IonOptiks) using an RSLC-nano uHPLC systems connected to a Lumos Fusion mass spectrometer (Thermo). Peptides were separated by a 70 min linear gradient from 5 % to 30 % v/v acetonitrile in 0.5 % v/v acetic acid. The mass spectrometer was operated in DIA mode, acquiring a MS 350-1650 Da at 120k resolution followed by MS/MS on 45 windows with 0.5 Da overlap (200-2000 Da) at 30 k with a NCE setting of 28. Ionised peptides were analysed in data-dependent mode and identified/quantified using DIA-NN version 1.8.2 beta 27 with Uniprot proteomes. Statistical analysis was performed using Wasim Aftab’s implementation of the Linear Models for Microarray Data (LIMMA) pipeline for two group comparison in a proteomic experiment ^89-91^

### TALE Design, Assembly and Transfection

TALE DNA binding domains targeting a 16 bp sequence proximal to the *Gata6* (TGGACTCGCTCCTACT), *Sox7* (TCCACAGTGCCAGTTT) and *Snai1* (TATCATCGCACTTTCT) TSSs were designed using the TAL Effector Nucleotide Targeter 2.0 software ^92^. TALEs were assembled by introducing four pre-assembled multimeric TALE repeat modules corresponding to these sequences into the TALE-VP64 backbone as described by Therizols et al (‘VP64’) using the approach described by Ding et al ^93,94^. A second set of control plasmids was created by cloning the same modules into a modified plasmid backbone in which we replaced the VP64 coding sequence with that encoding a non-functional 20 aa control peptide (‘Cntrl’; GGGSGGSGSMDAKSLTAWS).

ESCs were transfected with TALE plasmids using Lipofectamine 3000 Reagent according to manufacturer’s instructions (Life Technologies - L3000015). Briefly, A transfection mixture containing 1 μg of DNA (an equimolar pool of VP64 or control plasmids), 7.5 μl of Lipofectamine, 2 μl of P3000 reagent and Optimem (Thermo Fisher - 31985062) was combined with 0.5 × 10^6^ ESCs suspended in 1.25 ml of culture media supplemented with Pen/Strep. Each transfection was seeded in a gelatin coated well of a 6-well plate (see above) and cultured for 48 h, with a media change at 24 h. This was to provide sufficient time to ensure activation of their target genes. Cells were harvested by trypsinisation and checked for transfection efficiency by assessing the proportion of cells in the culture that express GFP (encoded in the plasmid backbone) measured using a NovoCyte flow cytometer (Agilent). A sample of d0/ESCs were retained to assess target gene activation and the remaining cells were differentiated for 14 days as described above. Transfections yielded on average > 70% GFP+ cells.

### RNA Isolation, NGS Library Preparation and Sequencing

RNA was extracted from 0.5 – 1 × 10^6^ cultured cells using Trizol reagent according to manufacturer’s instructions (Ambion - 15596026). RNA was DNase treated using TURBO DNase according to manufacturer’s instructions (Life Technologies - AM2238). RNA yield and integrity was determined using a combination of Qubit (Thermo Fisher), electrophoresis and TapeStation (Agilent) measurements. Strand-specific next generation sequence (NGS) library preparation was performed using the NEBNext rRNA Depletion Kit (NEB - E6310) in combination with the NEBNext Ultra II Directional RNA Library Prep Kit for Illumina (NEB - E7760) according to manufacturer’s instructions. Individual libraries were barcoded using NEBNext Singleplex primers (NEB - E7335, E7500, E7710 and E7730).

Pooled NSC libraries (**Fig. 1**) were sequenced as a 2 × 100 bp paired end run on the NextSeq 2000 platform (Illumina Inc - SY-415-1002) using P2 Reagents (200 cycles) v3 Kit (Illumina Inc - 20046812). Sequencing was performed by the Edinburgh Clinical Research Facility (ECRF). Pooled ES to NPC differentiation libraries (**Fig. 2**) were sequenced as a 2 × 51 bp paired end run on the novaseq_A00291 platform (Illumina) using the S1 Flow Cell and SCV3 chemistry (chemistry 1.5; revcomp index 2). Sequencing was performed by Edinburgh Genomics (EG). Pooled ES to NPC differentiation (TALE experiment; **Fig. 4**) and NSC (experiment 2; **Supplemental Fig. S4**) were sequenced as a 2 × 51 bp paired end run on the NextSeq 2000 platform (Illumina Inc - 20038897) using the P3 XLEAP-SBS Reagent Kit (100 cycles; Illumina - 20100990). Sequencing was performed by the ECRF.

### Chromatin Immunoprecipitation, NGS Library Preparation and Sequencing

#### Calibrated cross-linked ChIP (***Fig. 2***)

2 × 10^6^ NSCs were washed twice in PBS, resuspended in 250 µL of 1x PBS and fixed by the addition of an equal volume of 2x Formaldehyde Buffer (10 mM HEPES-KOH pH 7.9, 20 mM NaCl, 0.2 mM EDTA pH 8, 0.1 mM EGTA pH 8 and 2% methanol-free formaldehyde (Pierce - 28906)) and incubated for 10 min at RT. Fixation was quenched by incubation with 125 mM of glycine for 5 min at RT and then washed with 1x PBS. Fixed NSCs were combined with a 1/25^th^ of the number of equivalently fixed human HEK293 cells, snap frozen in liquid N2 and stored at -70 °C.

Cell pellets were thawed on wet ice, resuspended in ice cold Lysis Buffer 1 (50 mM HEPES-KOH pH 7.9, 140 mM NaCl, 1 mM EDTA pH 8, 10% v/v Glycerol, 0.5% v/v IGEPAL CA-630 (Merck - I8896) and 0.25% v/v Triton X-100) and incubated for 10 min at 4°C. Following centrifugation at 2000 x g for 5 min at 4 °C, the supernatant was discarded, and the pellet was resuspended in Lysis buffer 2 (10 mM Tris-HCl pH 8, 200 mM NaCl, 1 mM EDTA pH 8 and 0.5 mM EGTA) and incubated for 10 min at 4°C with rotation. Following centrifugation at 2000 x g for 5 min at 4 °C, the supernatant was discarded, and the pellet resuspended in Lysis buffer 3 (10 mM Tris-HCl pH 8, 100 mM NaCl, 1 mM EDTA pH 8, 0.5 mM EGTA, 0.1% w/v Na-Deoxycholate and 0.5% w/v N-Lauroylsarcosine). The nuclear extract was transferred to a 1 ml millitube with AFA Fibre (COVARIS - 520130) and sheared by sonication using an E220evolution Focused-ultrasonicator (COVARIS - 500429). Sonication was performed at 4 °C with an initial 30 s high intensity pulse (Peak Power = 75, Duty Factor = 20 and Cycle/Burst = 200) followed by a further 60 - 90 min of preparative sonication (Peak Power = 75, Duty Factor = 10 and Cycle/Burst = 200). Sonication conditions and timing were optimised to yield DNA fragments ranging from 200 - 500 bp. Chromatin was precleared by centrifugation at 16,000 x g for 10 min at 4 °C. The supernatant was transferred to a fresh tube and supplemented with BSA to a final concentration of 25 mg/ml. A 10 % sample of chromatin was retained as an input. Antibodies were adsorbed on to protein A Dynabeads (ThermoFisher - 10001D) by combining antibody and Dynabead suspension at a ratio of 1 μg to 30 μL and incubation for 1 h at 4°C. Chromatin from 1 x 10^6^ cells was combined with either 0.5 µg anti-H3K27me3 (C36B11; Cell Signalling – 9733S) or 0.5 µg anti-H2AK119ub (D27C4; Cell Signalling - 8240S), and incubated O/N on a rotating wheel at 4 °C. Bead-associated immune complexes were washed sequentially with ChIP Dilution Buffer, Wash Buffer A, and Wash Buffer B, each for 10 min at 4 °C on a rotating wheel, followed by two washes in TE buffer at RT (Wash Buffer A: 1% v/v Triton X-100, 0.1% w/v sodium-deoxycolate, 0.1% w/v SDS, 1mM EDTA, 500 mM NaCl and 50 mM HEPES at pH 7.9; Wash Buffer B: 0.5% v/v IGEPAL, 0.5% w/v sodium-deoxycolate, 1 mM EDTA, 250 mM LiCl and 20 mM Tris-HCl at pH 8.1). Purified DNA fragments were released by incubating the beads in 100 µL of elution buffer (0.1 M NaHCO3 and 1% w/v SDS) for 15 min at 37°C, followed by the addition of 50 µg of RNase A and 6 µL of 2 M Tris (pH 6.8) and incubation for 2 h at 65 °C and finally by the addition of 50 µg of proteinase K and incubation for 8 h at 65°C to degrade proteins and reverse the cross-links. The Dynabeads were removed using a magnetic rack and the isolated DNA was purified using PCR purification columns (Qiagen) according to manufacturer’s instructions. All buffers were filtered with a 40 µM filter and stored at 4 °C. NGS libraries were prepared using NEBNext Ultra II DNA Library Prep Kit for Illumina (NEB - E7103L). All lysis and wash buffers were supplemented with 1 mM DTT and 1× Protease inhibitors (Roche - 11836170001) just prior to use.

#### Non-calibrated cross-linked ChIP (***Fig. 3***)

Chromatin was prepared from 5 × 10^6^ cells and immunopurified with either anti-RING1B (D22F2; Cell Signalling - 5694) or anti-RYBP (Millipore - AB3637) as previously described, but without the inclusion of exogenous spiked in cells^23^. NGS libraries were prepared as previously described^95^.

#### Non-calibrated native ChIP (***Fig. 3***)

Chromatin was prepared from 1 – 3 × 10^6^ cells and immunopurified with either anti-H3K27me3 (C36B11; Cell Signalling – 9733S) or anti-H2AK119ub (D27C4; Cell Signalling - 8240S) as previously described^96^. NGS libraries were prepared as previously described^95^ with modifications outlined in ^96^.

#### Library Sequencing

NGS libraries, barcoded using NEBNext Singleplex primers (NEB - E7335, E7500, E7710 and E7730) were combined for sequencing as equimolar pools. Each pool was sequenced as a paired end (2 × 75 bp) run on a NextSeq 550 platform (Illumina Inc - SY-415-1002) using the High-Output v2.5 (150 cycle) Kit (Illumina Inc - 20024907). Sequencing was performed by the ECRF.

### Data Analysis

#### ChIP-seq mapping, processing and visualisation

Raw paired-end reads from calibrated ChIP-seq data were mapped to a custom merged index including both the mouse (mm39) and human (hg38) genomes using bowtie2 (v 2.4.2) ^97^ with the following parameters (-p 8 --local -D 20 -R 3 -N 1 -L 20 -i S,1,0.50 --no-mixed --no-discordant). Using the HOMER package (v4.10) ^98^, SAM files were converted into tag directories and multimapping reads were removed using makeTagDirectory with the following parameters (-unique -fragLength given -format sam). Mapped regions that, due to fragment processing, extended beyond the end of chromosomes were removed using removeOutOfBoundsReads.pl with chromosome lengths for mm39. Genome browser files (.bw) were generated using makeUCSCfile with the following parameters (bigWig -fsize 1e20-strand both -norm *norm_factor*). ChIP-seq signal was quantified at gene TSSs (Refseq mm39) using annotatePeaks.pl with the following parameters (-size "given" -noadj -strand both-noann). Quantification across the genome was performed using the same parameters using a custom interval (‘.bed’) file spanning the entire mouse genome (mm39) in 5 × 10^5^ bp windows with a 2.5 × 10^5^ bp slide/resolution. Heatmap quantifications for TSSs (Refseq mm39; ± 5 kb) were generated using annotatePeaks.pl with the following parameters (-size "given" -noann - nogene -noadj -hist 100 -ghist). All processed data was normalized to a calibration factor that was calculated based on the relative contribution of uniquely mapped human spike-in reads between the input and immunoprecipitated samples calculated as previously described ^99^.

Raw single-end reads from non-calibrated ChIP-seq were processed essentially as for calibrated ChIP-seq but were mapped to the mouse genome (mm39) alone. Genome browser files for all samples were normalised to 1 × 10^7^ uniquely mapped reads using the makeUCSCfile in the HOMER package.

All visualisations, plots, heatmaps and statistics were generated using custom R scripts. Heatmaps profiling calibrated ChIP-seq data were normalised based on the calibration factor and those for non-calibrated ChIP-seq were quantile normalised using the normalizeQuantiles function in the LIMMA package in R^90^.

#### RNA-seq mapping and analysis

Following demultiplexing, raw paired-end reads from RNA-seq data were aligned to the mouse genome (mm39 build) using the STAR aligner software (2.7.8a) ^100^ with default settings and --runThreadN 24. Aligned (‘.sam’) files were converted into tag directories using the HOMER package (v4.10) ^98^ using the makeTagDirectory function with the following parameters (-format sam -flip -sspe). Genomic intervals that extended beyond the end of the chromosomes were removed using the removeOutOfBoundsReads.pl function with chromosome lengths set to the mm39 genome build. Strand-specific browser track files (‘.bigWig’) were generated using the makeUCSCfile function with the following parameters (-fsize 1e20 -strand + or - -norm 5e6 (for NSCs or 10e6 for ESCs/NPCs)). The analyzeRepeats.pl function was used to quantify read coverage across all gene exons using the following parameters (none -count exons - strand + -noadj) based on a RefSeq annotation (mm39 genome build; ‘.gtf’ format). Using custom scripts in R, the quantified values were converted into reads per kilobase (RPK), with the addition of a nominal offset of 1 RPK. Differential gene expression analysis was performed using the LIMMA package for R/Bioconductor (http://www.R-project.org/)^90,101^. RPK values were read into R using the read.maimages function with source="generic", followed by normalisation across all samples using the normalizeBetweenArrays function with method=”quantile". Differential analysis by linear modelling, eBaysean statistics and multiple testing correction (Benjamini-Hochberg - ‘BH’) was performed for user defined experimental contrasts using the contrasts.fit, eBayes and topTable functions. Genes were identified as being differentially expressed if they had a corrected p-value of ≤ 0.01 and a log_2_ fold change (FC) of ≥ 1 or ≤ -1.

#### Motif and Functional Enrichment Analysis

Motif analysis was performed using the HOMER package (v4.10; ^98^) using the findMotifsGenome.pl function with -size given. Motifs enrichment was determined for the TSSs (RefSeq mm39; ± 500 bp) of upregulated genes (in I53A NSCs) with high H2AK119ub levels (top 25%) against a randomly sampled matched background (-bg) set of TSS with high H2AK119ub levels (top 25%) that were not differentially expressed.

Functional GO analysis was performed using the enrichGO function in the clusterProfiler R package with the following parameters (OrgDb = "org.Mm.eg.db", keyType = "SYMBOL", ont = "BP", pvalueCutoff = 0.05 and pAdjustMethod = "BH") and visualised using the ggplot2 package using custom R scripts ^102^.

## Data Availability

All high-level processed data is included as supplementary data. All raw next generation sequence (NGS) data is deposited in GEO (https://www.ncbi.nlm.nih.gov/geo/) under the following series accession (GSEXXXXXX). Computer codes and associated files for this study are available via GitHub (https://github.com/) in the ‘RIllingworth/Doyle_2025’ repository.

## ACKNOWLEDGEMENTS

Work in the Illingworth lab is supported by the MRC (Career Development award - MR/S007644/1) and the Simons Initiative for the Developing Brain (SIDB; SFARI - 529085). Work in the Adams group is supported by the MRC (MRC University Unit grants MC_UU_00007/6 and MC_UU_00035/3). L.A.D. and F.U.B. are supported by Doctoral Training Fellowships from Martin Lee and the Turkish Ministry of National Education respectively. We would like to thank Ms Aisling Fairweather for support and advice with NSC culture. We would like to thank all staff at the ECRF and Edinburgh Genomics for next generation sequencing and support. We would like to thank Dr Matthieu Vermeren for help with Microscopy and Image analysis, Ms Fiona Rossi and the IRR FACs facility for support with flow cytometry and the Edinburgh BioResearch and Veterinary Services for support with animal husbandry and research. Thank you also to the Lowell, Williams, Pollard and Emmerson labs for the kind gift of antibodies. We are grateful to Prof. Nick Gilbert, Prof. Wendy Bickmore, Prof. Steve Pollard, Prof. John Mason and members of the Illingworth lab for helpful Discussions and feedback on the manuscript.

## Figure Legends

**Supplementary Fig. 1: Characterisation of the developmental impact of *Ring1b* catalytic deficiency.**

**a,** Representative images of E12.5 embryos from a cross between *Ring1b^I53A/+^* heterozygote mice. The *Ring1B* genotype is indicated in each panel (scale bar = 1 mm).

**b,** Barplots comparing length measurements taken from *Ring1b^+/+^* (WT) and *Ring1b^I53A/I53A^*(I53A) E12.5 mouse embryos. Significant changes in crown to rump length, head width and forebrain width between WT and I53A embryos was determined using a Student’s Paired T-test, the results of which are displayed above each respective plot. Error bars represent the standard deviation of measurements taken from independent embryos (n=3 for crown to rump and n=5 for head and forebrain width for each genotype).

**c,** Representative bright-field images of *ex vivo* cultured primary telencephalic NSCs from *Ring1b^+/+^* (WT 1 & 2) and *Ring1b^I53A/I53A^* (I53A 1 & 2) mice. Images of two independently derived cultures are shown (scale bar = 100 μm). The right-hand panel shows a zoomed view as indicated.

**d,** Representative fluorescent images of DAPI staining in combination with either NESTIN or GFAP immunofluorescence in *ex vivo* cultured primary telencephalic NSCs derived from *Ring1b^I53A/I53A^* (I53A) E10.5 mouse embryos (scale bar = 100 μm; right hand panel shows a zoomed view as indicated). Images taken from a biologically independent I53A culture to that shown in Fig. 1c.

**e,** Genome browser tracks of normalised RNA-seq signal at candidate gene loci (*Sox2*, *Emx2*, *Sox9*, *Foxg1*, *Twist1* and *Tbx15*) from three independent *Ring1b^+/+^* (WT1-3) and *Ring1b^I53A/I53A^*(I53A1-3) NSC cultures. The signal is coloured according to the transcribed strand (positive - black and negative - grey) and annotated as per the mm39 genome assembly.

**Supplementary Fig. 2: Chromatin profiling in *Ring1b^+/+^* and *Ring1b^I53A/I53A^* ex vivo cultured NSCs.**

**a,** Genome browser tracks depicting calibrated ChIP-seq profiles for H3K27me3 (red) and H2AK119ub (blue) at candidate genomic loci (*Hoxd* cluster, *Esrrb* and *Tbx15*) in the indicated NSC cultures. The regions are annotated as per the mm39 genome assembly and read-depth ranges are noted for each immunoprecipitation.

**b,** Boxplots comparing calibrated ChIP-seq signal for H2AK119Ub (log_2_ FC *Ring1b^I53A/I53A^*/*Ring1b^+/+^*) at TSS (±2.5 kb) for the indicated NSC cultures. Significant changes between datasets (determined using a Wilcoxon rank sum test) are indicated above their respective plot.

**c,** Scatter plots showing the H2AK119ub1 levels at TSSs (±2.5 kb) in the indicated NSC cultures. H2AK119ub values for TSSs of genes that are upregulated in *Ring1b^I53A/I53A^* NSCs are highlighted in red.

**d,** Heatmaps of H3K27me3 ChIP-seq signal spanning ±5 kb of all gene TSSs (left) and the top enriched 10% of TSS (right) in the indicated NSC cultures. Heatmaps are ranked from high to low signal based on the WT1 NSC dataset (mean of two replicate experiments). Summary profiles of the mean calibrated ChIP-seq signal are presented for each heatmap for the indicated NSC cultures.

**e,** Heatmaps of H3K27me3 calibrated ChIP-seq signal across the TSSs (±5 kb) of genes upregulated in *Ring1b^I53A/I53A^* (I53A) NSCs.

**f,** Scatter plots comparing differential absolute levels of H2AK119ub and H3K27me3 in the indicated NSC cultures and bounded by summary boxplots for each calibrated ChIP-seq dataset. Differential signal represents calibrated ChIP-seq signal at TSSs (±2.5 kb) comparing *Ring1b^I53A/I53A^* (I53A) to *Ring1b^+/+^* (WT) NSCs. Pearson correlations for all (grey) and for genes upregulated in *Ring1b^I53A/I53A^* NSCs (red) are shown in parenthesis. Significant changes between datasets (determined using a Wilcoxon rank sum test) are indicated next to their respective plot.

**g,** TF Motifs (HOMER de novo motif screen – see methods) found to be significantly enriched in the TSSs (±500 bp – see cartoon schematic) of genes that are upregulated in *Ring1b^I53A/I53A^*vs. *Ring1b^+/+^* NSCs. Heatmap showing RNA levels for the genes encoding the TFs (normalised RPK values scaled as per the colour bar). Filled circles indicate detectable protein expression in *Ring1b^+/+^* NSCs as measured by Mass Spectrometry.

**Supplementary Fig. 3: Gene expression and chromatin profiling in *Ring1b^+/+^* and *Ring1b^I53A/I53A^* cells during in vitro NPC differentiation.**

**a,** Schematic of ES to NPC differentiation and sample collection points.

**b,** Representative bright-field images of *Ring1b^+/+^* and *Ring1b^I53A/I53A^* cells at day 0 and 14 of NPC differentiation (Independent replicate from that shown in Fig. 3b; Scale bars represent 100 μm).

**c,** Representative fluorescent images of DAPI staining and TUJ1 (β-tubulin) immunofluorescence from two independent NPC differentiations. Shown are images for *Ring1b^+/+^* and *Ring1b^I53A/I53A^* cells at day 14 of differentiation (Independent replicates from those shown in Fig. 3c; Scale bars represent 100 μm).

**d,** Scatter plots comparing log_2_ mean expression vs. log_2_ fold changes (FC; log2 Day X/Day 0) for *Ring1b^+/+^*and *Ring1b^I53A/I53A^* cells at day 3, 7, 10 and 14 of NPC differentiation. Differentially expressed genes (DEGs) which are significantly up- or downregulated are indicated in red and blue respectively (log_2_ FC ≥ 1 / ≤ -1 and p ≤ 0.01). Candidate marker genes and the total number of DEGs are indicated for each comparison.

**e,** Genome browser tracks of normalised RNA-seq signal at the *Gata4* locus at day 0, 3, 7, 10 and 14 of NPC differentiation for *Ring1b^+/+^* (upper) and *Ring1b^I53A/I53A^* (lower) cells. The signal is coloured according to the transcribed strand (positive - black and negative - grey) and annotated as per the mm39 genome assembly. Normalised read-depth range is indicated.

**f,** Scatter plots comparing log_2_ mean expression vs. log_2_ FC (I53A/WT) at day 0, 3, 7, 10 and 14 of NPC differentiation. Differentially expressed genes (DEGs) which are significantly up- or downregulated are indicated in red and blue respectively (log_2_ FC ≥ 1 / ≤ -1 and p ≤ 0.01). Candidate marker genes and the total number of DEGs are indicated for each comparison.

**g and h,** Heatmaps depicting the log_2_ FC (log_2_ I53A/WT) for all upregulated **g**) and downregulated **h**) genes with their combined meta-profiles (dashed line = median, solid line = mean, dark grey = 25^th^ -75^th^ percentile and light grey 10^th^ – 90^th^ percentile). The heatmap is scaled as per the axis colour bar and a selection of candidate genes are shown for reference. **i,** Genome browser tracks of ChIP-seq signal for H3K27me3, H2AK119ub, RYBP and RING1B at the *Gata4* locus in WT ESCs (day 0) and their derivative NPCs (day 7). The region is annotated as per the mm39 genome assembly and read-depth range is indicated.

**j,** The top five enriched functional gene ontology terms for downregulated DEGs and heatmap of log_2_ FC (I53A/WT) for the indicated functional gene list. The heatmaps and profiles are shown as for **g** and **h**.

**Supplementary Fig. 4: Downregulation of neurogenic fate genes is a common feature of both ex vivo and in vitro derived *Ring1b^I53A/I53A^* NSCs.**

**a,** Representative bright-field images of in vitro derived NSCs from *Ring1b^+/+^* and *Ring1b^I53A/I53A^* cells (scale bar = 100 μm).

**b,** Dendrogram comparing gene expression profiles (normalised RNA-seq RPK values) in ES to NPC differentiating samples, in vitro derived NSCs and ex vivo derived NSCs (greyscale, blue and red respectively). Samples were ordered based on unsupervised hierarchical clustering.

**c,** PCA plots of the expression profiles shown in **b**, for PC1 v PC2 and PC2 v PC3 (left and right plots respectively). Points are coloured as per the colour bar in **b**, the dashed line indicates the ES to NPC differentiation trajectory and the relative variance per PC is indicated in parenthesis.

**d,** Heatmaps depicting the z-score values for candidate marker gene sets comparing ESCs (DO/ESC), NPCs (InV NPC; day 14), in vitro derived NSCs (InV NSC) and ex vivo derived NSCs (ExV NSC). The heatmap is scaled as per the colour bar.

**e,** Scatter plots comparing log_2_ mean expression vs. log_2_ fold changes (FC; log_2_ I53A/WT) for in vitro derived NSCs. Differentially expressed genes (DEGs) which are significantly up- or downregulated are indicated in red and blue respectively (log_2_ FC ≥ 1 / ≤ -1 and p ≤ 0.01). The total number of DEGs are indicated for each comparison.

**f,** Barplot showing the total number of DEGs (up and down regulated shown in red and blue respectively) for in vitro derived, individual ex vivo and combined ex vivo NSCs.

**g,** Venn diagram showing the number of DEGs in in vitro and ex vivo derived NSCs and the intersect between the two sets (up and down regulated; upper and lower panel respectively). The significance of the overlap for both sets of genes was determined using a Fisher’s Test, the result of which is shown below each respective Venn.

**h,** The top ten ontology terms enriched in downregulated DEGs (I53A vs WT) in in vitro and ex vivo derived NSCs. Colour indicates the adjusted p value and the circle size indicates the relative gene count for that GO term.

**Supplementary Fig. 5: Pronounced alterations to the gene expression profile of in vitro differentiated NPCs following ectopic TF activation.**

**a,** Representative brightfield images of un-transfected WT (WT UT), WT transfected with TALE-delta control (WT Cntrl), WT transfected with TALE-VP64 (WT VP64) and un-transfected I53A (I53A UT) ESCs (scale bars = 100 μm).

**b,** Representative brightfield images of cells treated as in panel **a** and cultured for 14 days under NPC differentiation conditions. Images represent an independent experiment from that shown in Fig 4c. Open and filled arrows indicate neurite projections and cobble-stone-like cells respectively (scale bars = 100 μm).

**c,** Scatter plots of log_2_ mean gene expression vs. log_2_ fold change (FC) for WT cells for the indicated comparisons at day 0 (ESCs; upper) and 14 (NPCs; lower) of differentiation. Differentially expressed genes (DEGs) that are significantly up- or downregulated are indicated in red and blue respectively (log_2_ FC ≥ 1 / ≤ -1 and p ≤ 0.01). Candidate marker genes and the total number of DEGs are indicated for each comparison.

**d,** Genome browser tracks of mean normalised RNA-seq signal at candidate gene loci (*Myl3* and *Gata4*) in ESCs and NPCs (day 14) for the indicated conditions. The signal represents the sum of three independent experimental replicates and is coloured according to the transcribed strand (positive - black and negative - grey). Regions are annotated as per the mm39 genome assembly.

Supplementary Table 1. Primary Cultured NSCs - Expression Analysis

Supplementary Table 2. Primary Cultured NSCs - Gene Ontology Analysis Supplementary Table 3. Primary Cultured NSCs - ChIP Analysis

Supplementary Table 4. Primary Cultured NSCs - Mass Spectrometry Analysis Supplementary Table 5. Primary Cultured NSCs - Motif Enrichment Analysis Supplementary Table 6. NPC Differentiation - Expression Analysis

Supplementary Table 7. NPC Differentiation - ChIP Analysis

Supplementary Table 8. NPC Differentiation - Gene Ontology Analysis Supplementary Table 9. In vitro vs. Ex Vivo NSCs - Expression Analysis

Supplementary Table 10. In vitro vs. Ex Vivo NSCs - Gene Ontology Analysis Supplementary Table 11. NPC Differentiation (TALE) - Expression Analysis

