## Supplementary figures and images for "H2AK119ub Safeguards Against Ectopic Transcription Factor Mediated Gene Activation in the Developing Forebrain"

### Supplementary Figures

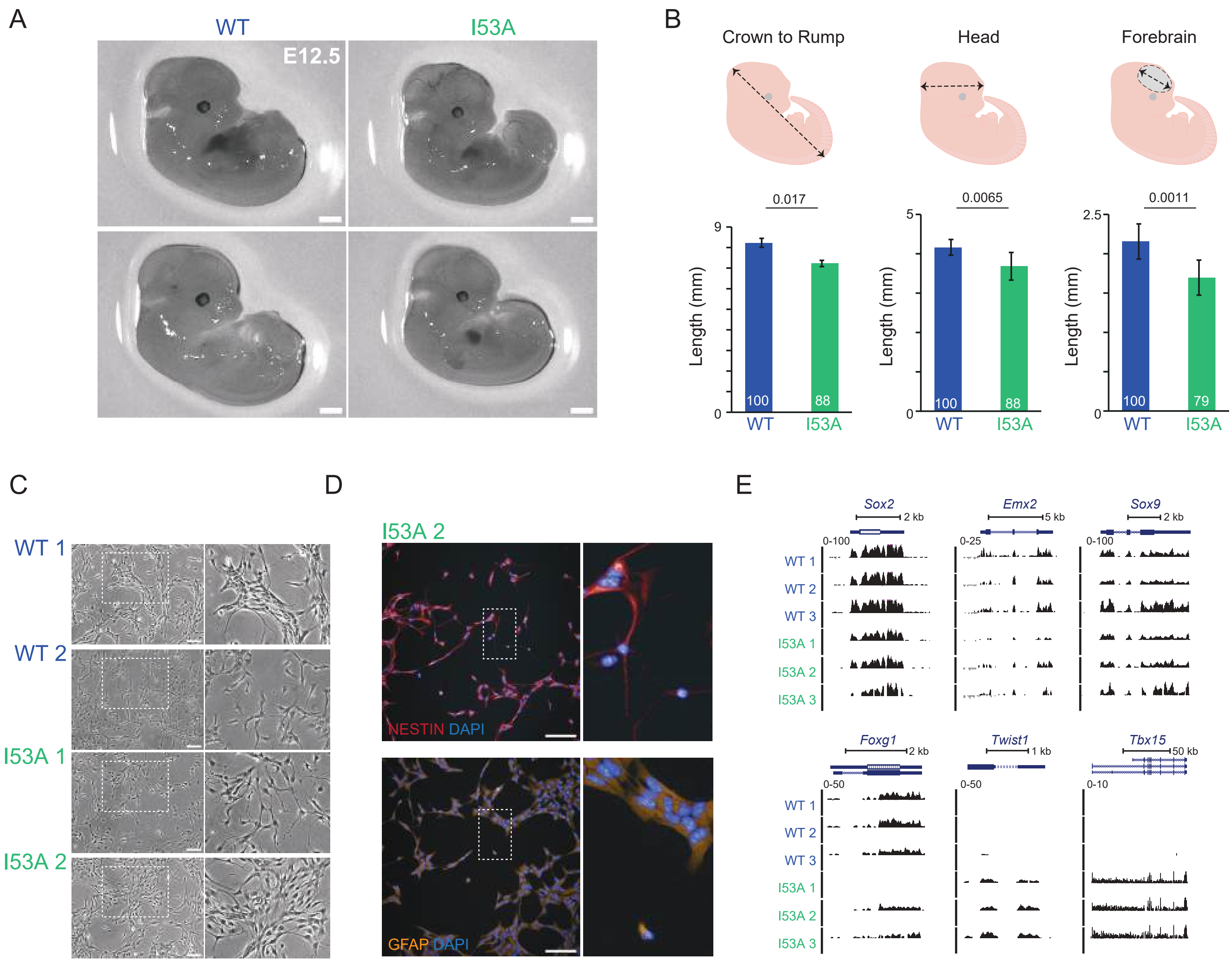

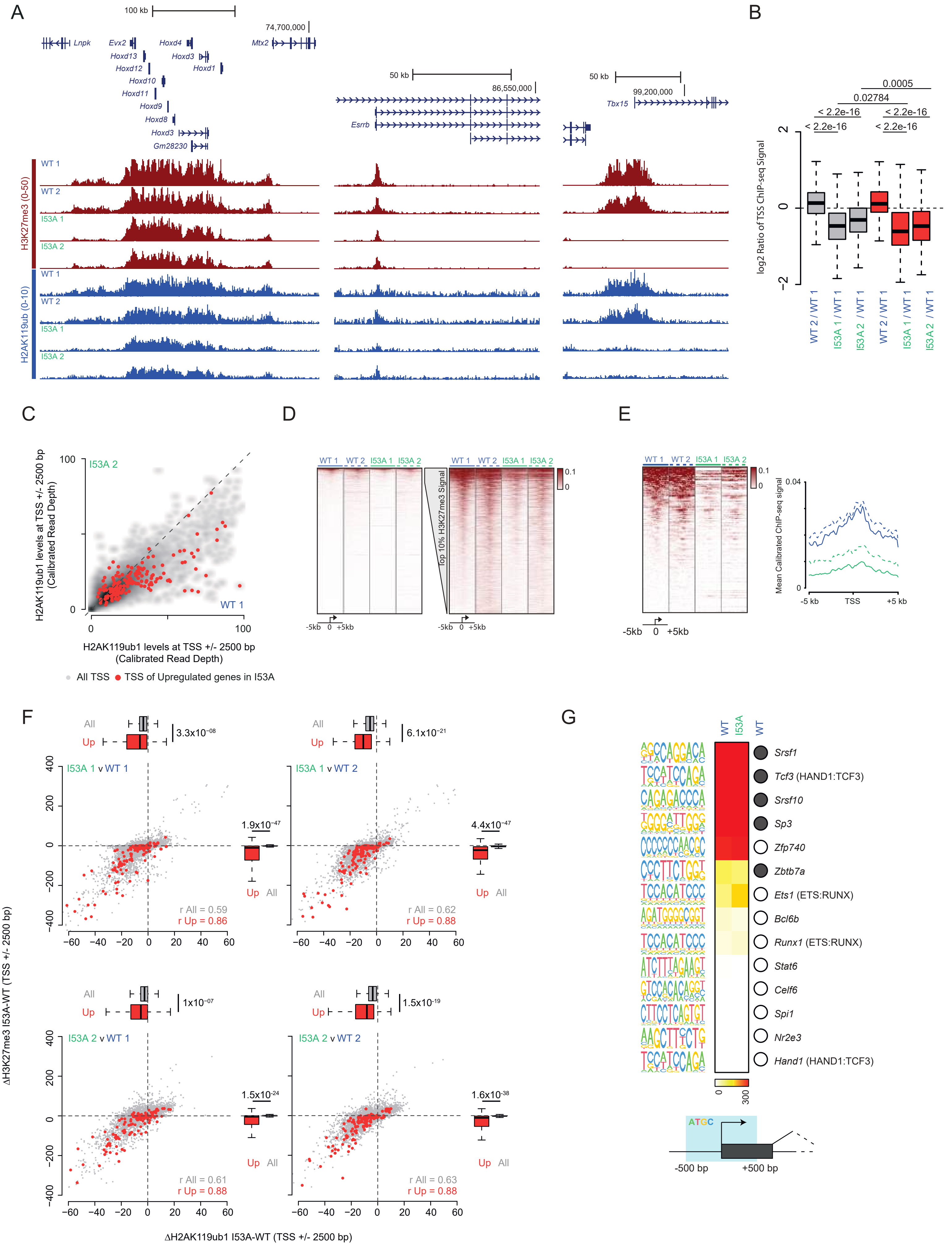

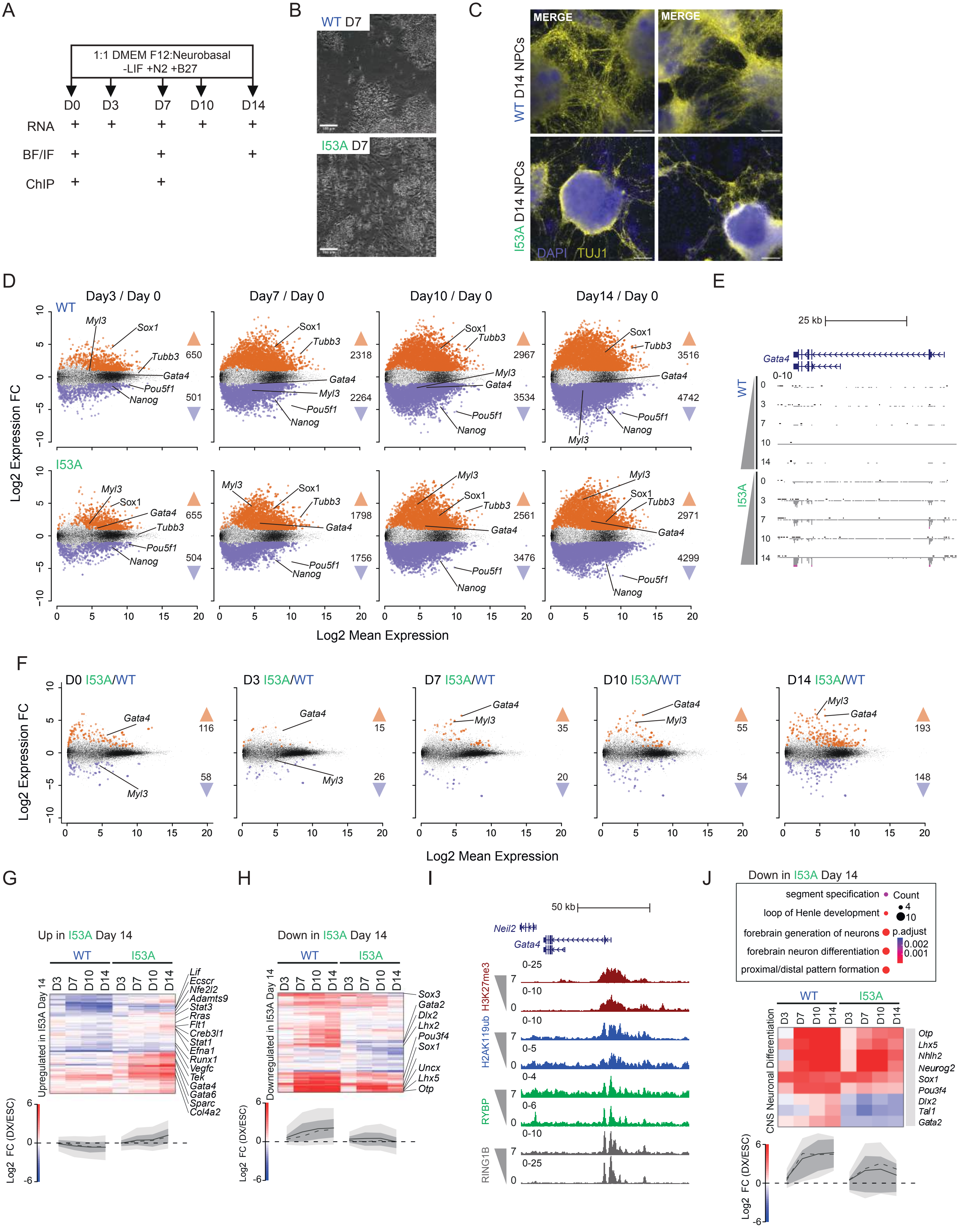

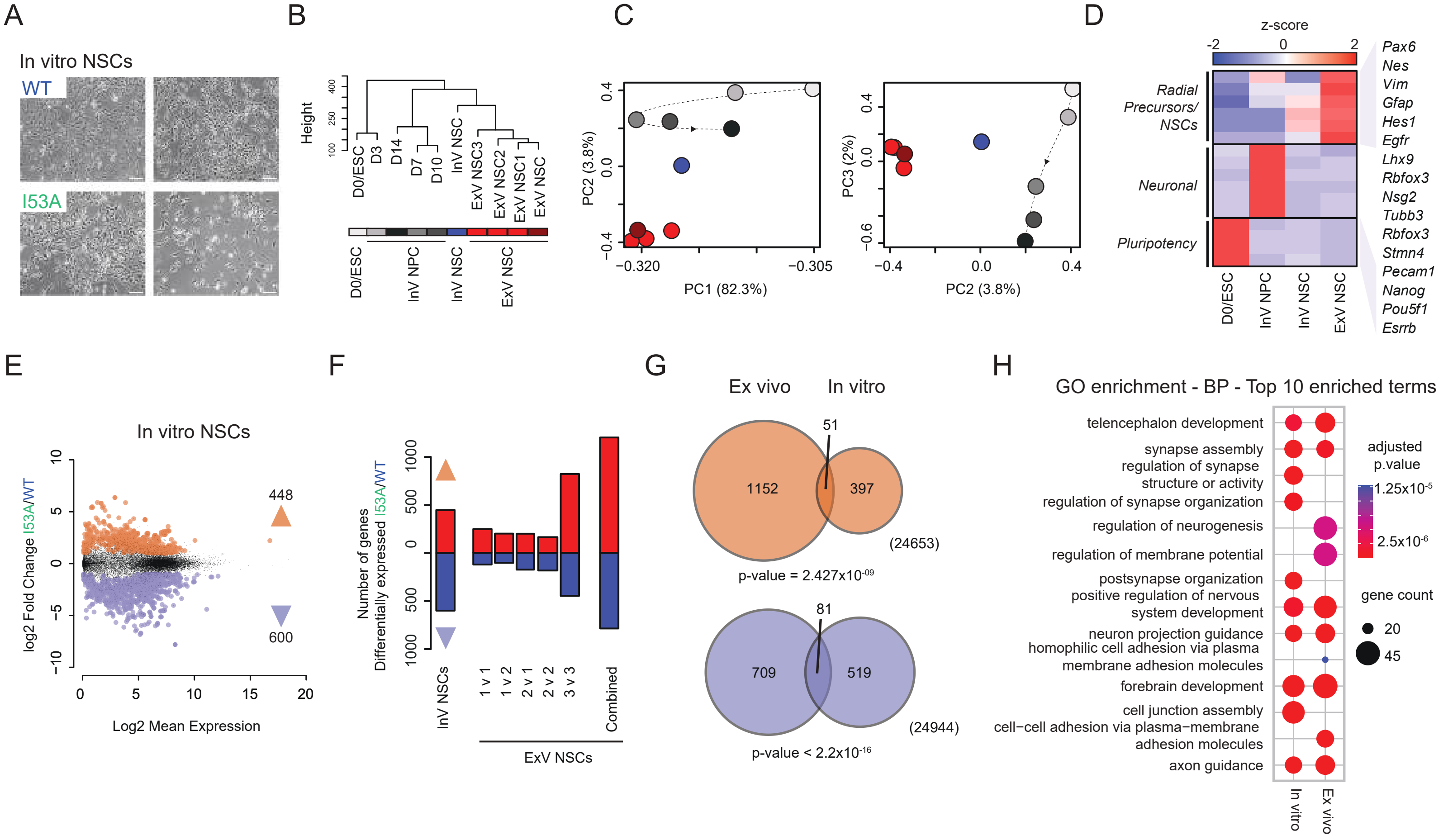

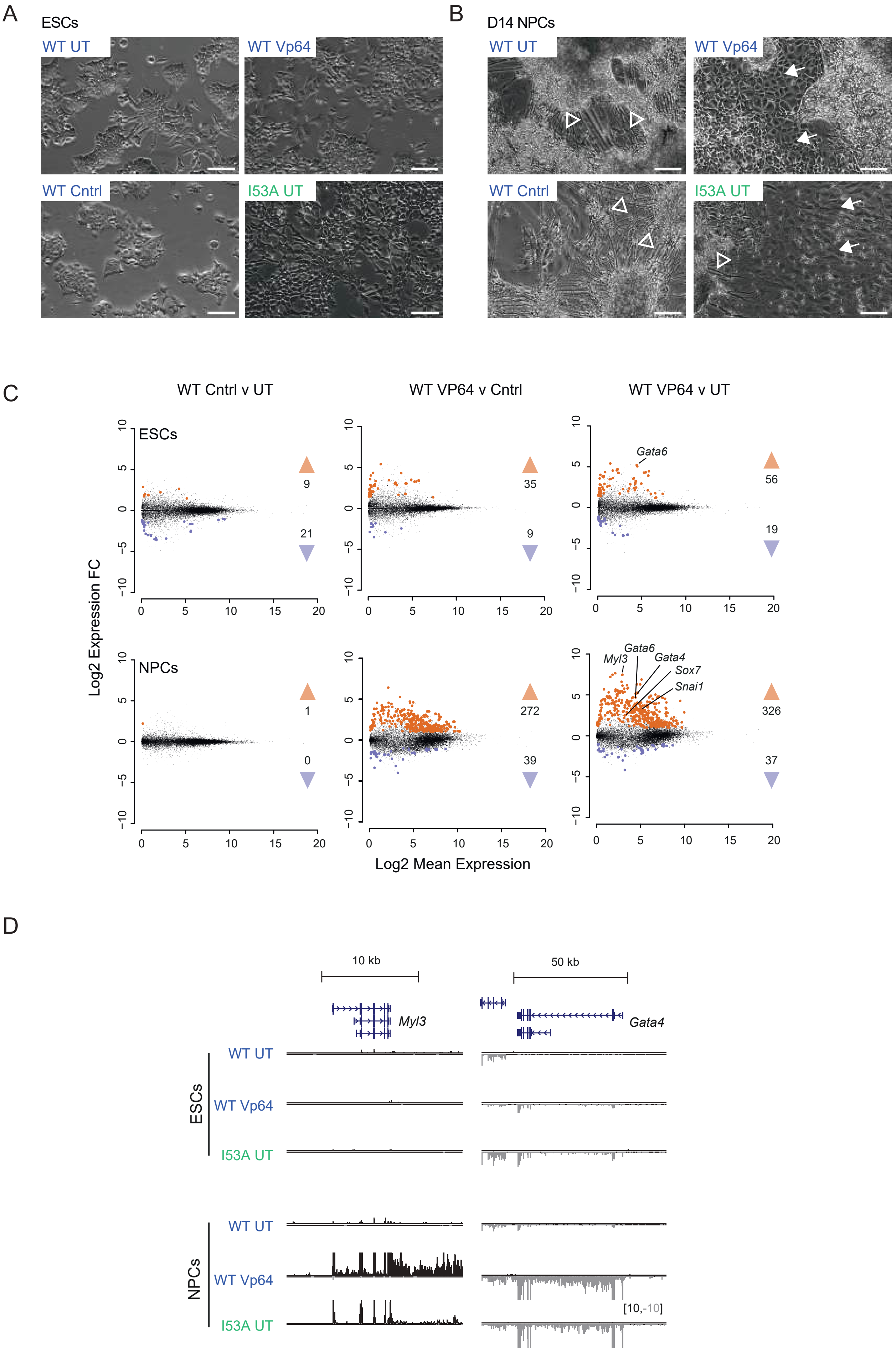
